# Maximizing binary interactome mapping with a minimal number of assays

**DOI:** 10.1101/530790

**Authors:** Soon Gang Choi, Julien Olivet, Patricia Cassonnet, Pierre-Olivier Vidalain, Katja Luck, Luke Lambourne, Kerstin Spirohn, Irma Lemmens, Mélanie Dos Santos, Caroline Demeret, Louis Jones, Sudharshan Rangarajan, Wenting Bian, Eloi P. Coutant, Yves L. Janin, Sylvie van der Werf, Philipp Trepte, Erich E. Wanker, Javier De Las Rivas, Jan Tavernier, Jean-Claude Twizere, Tong Hao, David E. Hill, Marc Vidal, Michael A. Calderwood, Yves Jacob

## Abstract

Complementary assays are required to comprehensively map complex biological entities such as genomes, proteomes and interactome networks. However, how various assays can be optimally combined to approach completeness while maintaining high precision often remains unclear. Here, we propose a framework for binary protein-protein interaction (PPI) mapping based on optimally combining assays and/or assay versions to maximize detection of true positive interactions, while avoiding detection of random protein pairs. We have engineered a novel NanoLuc two-hybrid (N2H) system that integrates 12 different versions, differing by protein expression systems and tagging configurations. The resulting union of N2H versions recovers as many PPIs as 10 distinct assays combined. Thus, to further improve PPI mapping, developing alternative versions of existing assays might be as productive as designing completely new assays. Our findings should be applicable to systematic mapping of other biological landscapes.

---

Complex biological entities such as genomes, proteomes and interactome networks require multiple, orthogonal approaches to be fully characterized. Even in the case of DNA, a highly homogenous macromolecule, sequencing and assembly of large genomes such as the human genome requires combinations of complementary methodologies including short-read sequencing as those provided by Illumina platforms, long-read DNA sequencing as in PacBio and/or Nanopore technologies, and HiC-related chromatin mapping tools^1^.

Because polypeptides are more heterogeneous and structurally complex macromolecules than DNA^2^, determining protein features at the scale of the whole proteome is considerably more challenging than characterizing nucleic acid sequences. For instance, to expand the detectable portion of a proteome by mass-spectrometry, negative peptide ionization approaches are necessary to complement canonical positive-mode strategies^3^. Another example is protein crystallization that often requires hundreds of different conditions to be tested, including combining parameters such as ionic strength, temperature, pH, salt, protein concentrations and others^4^. Structural proteomics, which determines protein structures at proteome-scale^5^ also requires a palette of technologies such as X-ray crystallography, nuclear magnetic resonance (NMR), and/or cryogenic electron microscopy (cryo-EM) to reasonably cover proteins and protein complexes of different sizes, solubilities, and flexibilities^6, 7^.

In short, the mapping of large macromolecular landscapes usually requires multiple technologies that can be combined to complement each other’s relative shortcomings. However, for most comprehensive mapping projects, it remains unknown how to optimally combine various experimental methods in order to achieve maximal detection and specificity by a minimal number of assays.

Macromolecules including DNA, RNA and proteins do not work in isolation but assemble into molecular machines, signaling cascades and metabolic pathways through interactions with other macromolecules, altogether forming complex interactome networks that are critical for biological systems^8, 9^. As protein-protein interactions (PPIs) play a pivotal role in these networks, it is essential to comprehensively identify binary PPIs^10–12^ and co-complex protein associations^13–15^, both at the scale of signaling pathways^16, 17^, molecular machines^18, 19^ and network modules^20^, and at the scale of the whole proteome^12^.

Although medium-to high-throughput binary PPI detection assays have been available for three decades, since the original description of the yeast two-hybrid (Y2H) system^21^, it remains impossible to completely map interactome networks using any assay in isolation. This is well illustrated by the fact that most binary PPI assays developed so far only detect about a third of well-described benchmark interactions from a positive reference set (PRS) under conditions that limit, but not necessarily completely prevent, the detection of negative control protein pairs from a random reference set (RRS)^22–27^. However, since different assays^23^ or assay versions^28, 29^ often detect overlapping, yet distinct subsets of interactions, combining them should significantly increase overall detection.

In order to comprehensively map binary PPIs both at the scale of signaling pathways and protein complexes and at the scale of whole proteomes, it should thus be possible to establish an optimal combination of assays and/or assay versions to capture nearly all interactions in a defined search space while ensuring high specificity (**Fig. 1a**). Our goal here was to systematically analyze experimental variables that influence binary PPI detection across multiple assays and integrate such knowledge into the rational design of a framework that maximizes detection with a minimal number of assays.

**Figure 1.**
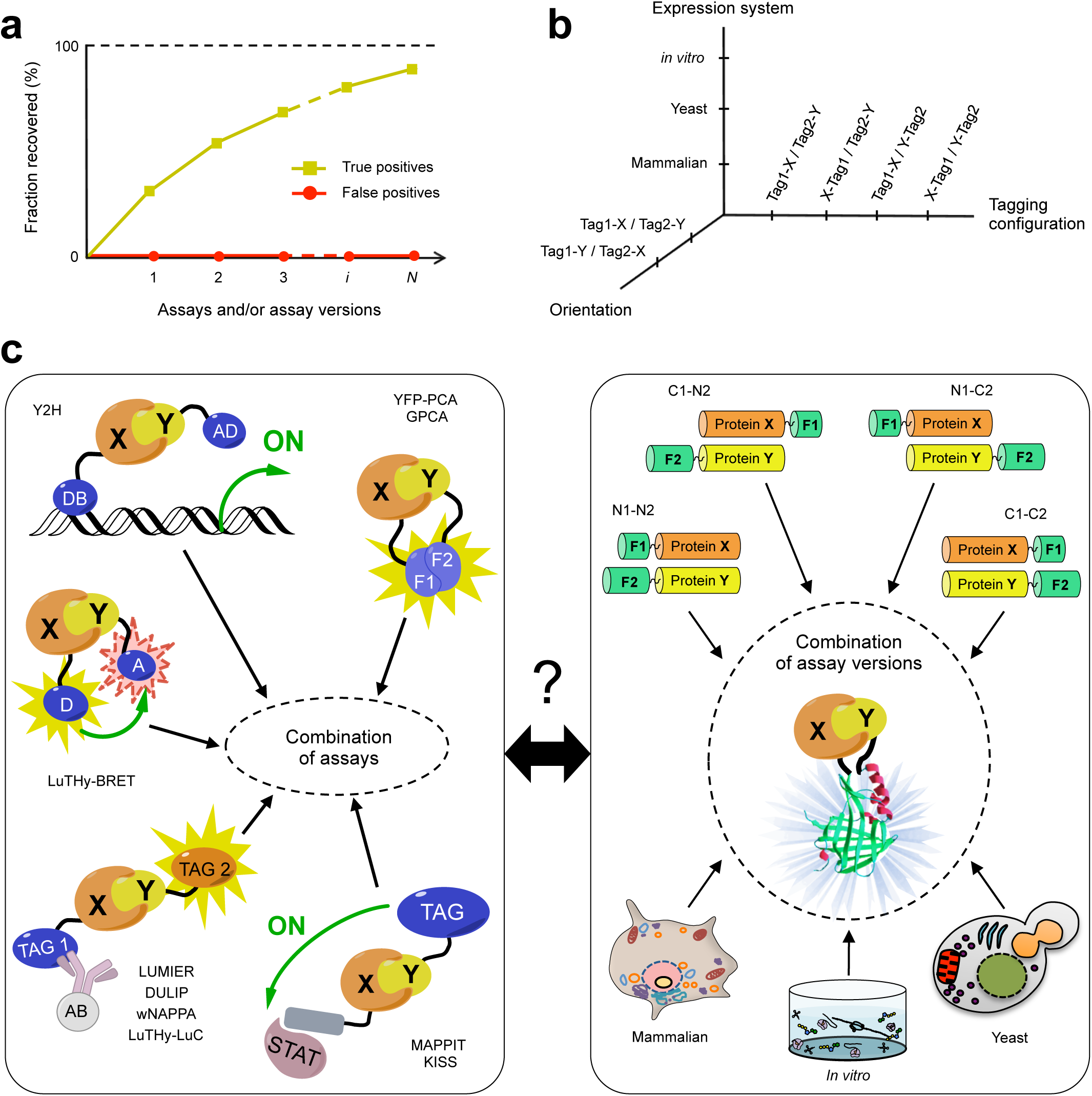
A framework to combine assays and maximize overall detection. (**a**) Combining multiple assays and/or assay versions to increase detection under maximum specificity. (**b**) A multi-dimensional space of experimental conditions for each binary PPI assay. (**c**) Comparing combinations of different, distinct assays (left) to combinations of multiple assay versions explored by the single, highly versatile NanoLuc 2-hybrid (N2H) system (right).

We evaluated a class of binary PPI assays that are based on exogenously expressing pairs of hybrid proteins, collectively referred to as Tag1-X and Tag2-Y, where X and Y can be any protein, and Tag1 and Tag2 are two polypeptidic moieties, the physical proximity of which can be readily monitored. For example, in the Y2H system, X and Y interacting proteins restore physical proximity between the DNA binding domain (DB) and the activation domain (AD) of a transcription factor upon exogenous expression of DB-X and AD-Y hybrid proteins^21, 30^.

In addition to assessing different assays defined by specific tag pairs, we also evaluated assay version differences due to: i) the relative orientation or swap of the tested proteins X and Y, differentiating between Tag1-X/Tag2-Y and Tag1-Y/Tag2-X; ii) the tagging configurations where we distinguish protein fusions from either the N-or C-terminal extremities of X and Y, *e.g.* Tag1-X/Tag2-Y versus X-Tag1/Y-Tag2; and iii) the exogenous expression systems, either in yeast, in mammalian cells, or *in vitro* (cell-free) (**Fig. 1b**).

To uniformly compare these different experimental conditions within a single assay, we have engineered a PPI detection technology based on the functional reconstitution of NanoLuc, a small 19kDa, high-performance bioluminescent protein^31^. Using this NanoLuc two-hybrid (N2H) system in combination with highly versatile vectors capable of driving protein expression as N-or C-terminal fusions of assay tags in all three environments, we show that permuting various experimental conditions in a single assay can achieve similar PPI detection as through the continuous addition of fundamentally different techniques (**Fig. 1c**). Our results suggest that only a few versatile assays should be sufficient to achieve near saturation with maximal precision in medium-and high-throughput proteome-scale PPI mapping projects. This finding might also be of value for large-scale mapping of other complex biological landscapes.

## RESULTS

### Combining binary PPI assays under maximized specificity

Different binary PPI assays often detect distinct subsets of interactions^23^, thus combining several assays should increase the overall recovery in a given search space of the proteome (**Fig. 1a**). However, what remains unclear is how binary PPI assay combinations affect the overall specificity, *i.e.* the fraction of true interactions detected in a given search space over the fraction of spurious pairs reported.

Ten years ago, our group introduced an empirical framework to benchmark binary PPI assays based on: i) a list of ∼100 well-documented positive control PPIs, constituting the first version of a PRS, or hsPRS-v1, ii) a list of ∼100 pairs of proteins randomly picked among the ∼2×10^8^ pairwise combinations of human proteins, constituting version 1 of a RRS, or hsRRS-v1^22^; and iii) the Gateway cloning of all open reading frames (ORFs) involved, allowing identical constructs to be used across many different assays^33^. We and others have since cross-examined the following binary PPI assays against each other: i) Y2H^23^, ii) yellow fluorescent protein-protein complementation assay (YFP-PCA)^23^, iii) mammalian protein-protein interaction trap (MAPPIT)^23^, iv) well-based nucleic acid programmable protein arrays (wNAPPA)^23^, v) luminescence-based mammalian interactome (LUMIER)^23^, vi) kinase substrate sensor (KISS)^25^, vii) *Gaussia princeps* complementation assay (GPCA)^24, 34^, viii) dual luminescence-based co-immunoprecipitation (DULIP)^26^, and ix) bioluminescence-based two-hybrid (LuTHy)^27^, combining bioluminescence resonance energy transfer (LuTHy-BRET) and luminescence-based co-precipitation (LuTHy-LuC) assays (**Fig. 1c**, left panel).

Under the various conditions used in these studies, each of the corresponding 10 assays, representing the union of only one or two version(s) (**Supplementary Table 1**), recovered between 21% and 39% of hsPRS-v1 pairs, with the negative control hsRRS-v1 detection ranging between 0% and 4% (**Supplementary Fig. 1a**). However, to directly compare the different assays side-by-side and combine them homogeneously, an identical RRS threshold should be applied to each individual assay. After applying an identical RRS cutoff, increasing from 1% to 10%, to each of the above-mentioned assays, we observed that the union of the ten different assays detected between 70% and 91% of positive control hsPRS-v1 pairs with 8% and 59% of random hsRRS-v1 pairs scoring positive, respectively (**Supplementary Fig. 1b**). Therefore, to avoid accumulation of random pairs affecting the quality of future analyses, each assay should be calibrated such that none of the RRS pairs are scored positive. After reanalyzing published data under these conditions, the recovered fractions of hsPRS-v1 pairs ranged from 3% to 33% for individual assays performed in only one or two version(s) (**Fig. 2a**), while the cumulative detection reached 65% when all assays were combined (**Fig. 2b** and **Supplementary Fig. 1c**). This suggested that many different assays, each associated to specific tag pairs and plasmids, would be needed to reach full coverage.

**Figure 2.**
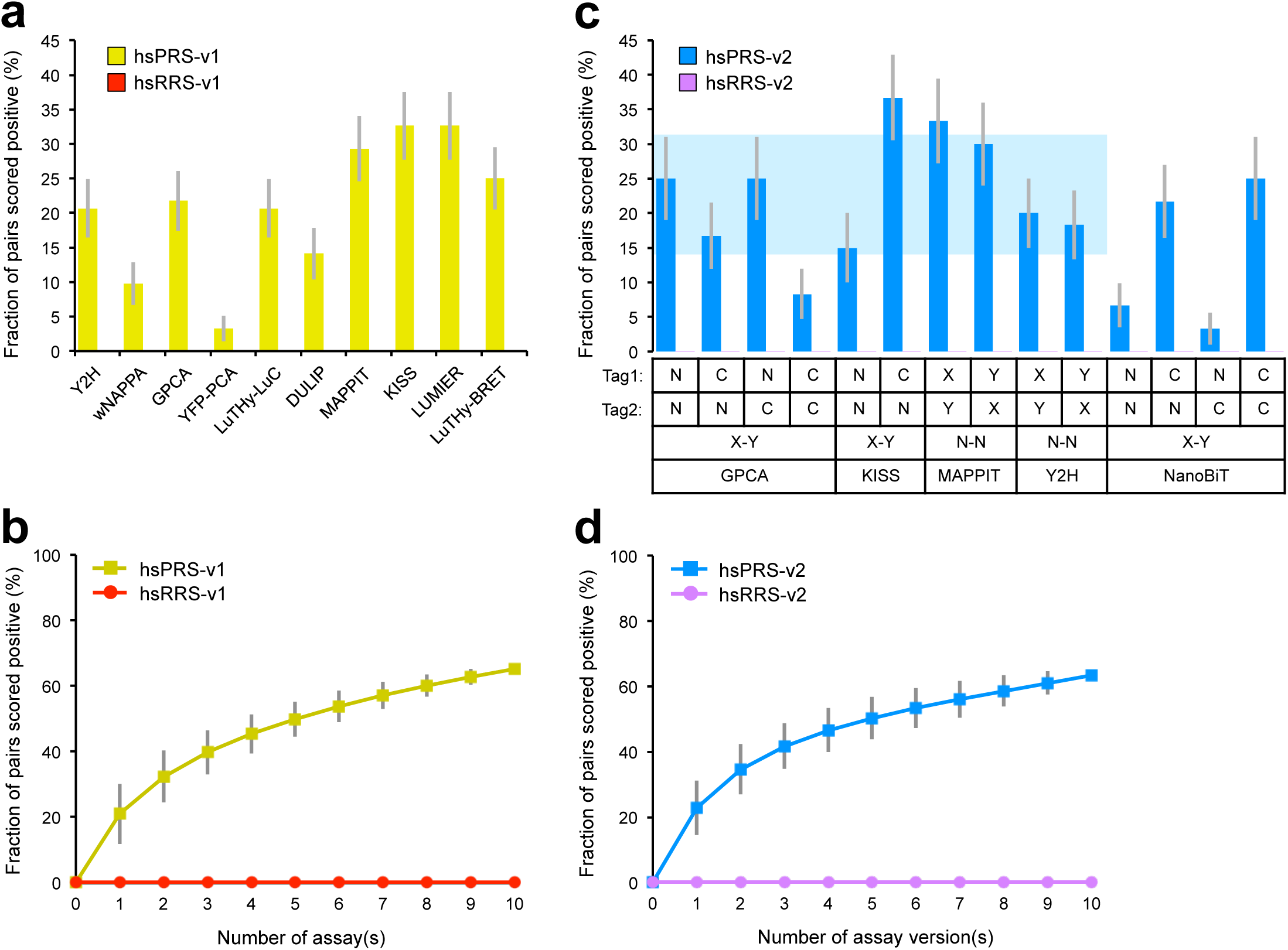
Recovery of positive control PRS pairs by multiple assays or assay versions. (**a**) Fractions of hsPRS-v1 pairs recovered by ten well-established assays when an identical threshold of no hsRRS-v1 pairs scoring positive is applied. (**b**) Cumulative hsPRS-v1 recovery rates when assays in (**a**) are combined (individual data points are displayed in **Supplementary Fig. 1c**). (**c**) Fractions of hsPRS-v2 pairs recovered by different versions of GPCA, KISS, MAPPIT, Y2H and NanoBiT when an identical threshold of no hsRRS-v2 pairs scoring positive is applied. Horizontal shaded blue area indicates the average recovery rate ± S.D. for GPCA, KISS, MAPPIT and Y2H versions. (**d**) Cumulative hsPRS-v2 recovery rates when versions of GPCA, KISS, MAPPIT and Y2H presented in (**c**) are combined (individual data points are displayed in **Supplementary Fig. 2d**). Error bars indicate standard errors of proportions (**a**,**c**) or standard deviations (**b,d**).

### Benchmarking binary PPI assays and assay versions

We considered the possibility that the failure of the 10 assays described above (**Fig. 2a,b**) to detect nearly all PRS interactions might be due to ORF clone quality issues and/or problematic interaction annotations. To avoid such possibilities, we re-examined all protein pairs from the original reference sets: hsPRS-v1 and hsRRS-v1^23^. As a first step, only fully sequence-verified, full length clonal ORFs were isolated (**Supplementary Fig. 2a**). Resulting pairs were then filtered based on updated literature information and annotations for binary PPIs (**Supplementary Table 2**). Finally, 60 PRS and 78 RRS pairs were assembled into a second-generation of reference sets: hsPRS-v2 and hsRRS-v2, respectively (see **Methods** for details and **Supplementary Fig. 2b**). After repeating four assays that were originally benchmarked against hsPRS-v1 and hsRRS-v1 with the second generation of reference sets, the cumulative PRS recovery rate reached 50% and 58% with -v1 and -v2 sets respectively, when none of the RRS pairs were scored positive (**Supplementary Fig. 2c**).

As the failure of 10 well-established assays to detect nearly all binary PPIs did not arise from problems with the original reference sets, we decided to evaluate whether expanding the number of experimental conditions, or assay versions (**Fig. 1b**), covered by any single technique could increase its coverage and ultimately reduce the number of assays required. To investigate this, we benchmarked several versions of Y2H, MAPPIT, KISS and GPCA against the second-generation of reference sets, hsPRS-v2 and hsRRS-v2. These four assays and corresponding versions (10 in total) were implemented as summarized in **Supplementary Table 1**. In particular, we constructed Gateway-compatible vectors for GPCA to cover all four possibilities of tagging configurations with this assay, *i.e.* Tag1-X/Tag2-Y, X-Tag1/Tag2-Y, Tag1-X/Y-Tag2 and X-Tag1/Y-Tag2 or, simply, N1N2, C1N2, N1C2 and C1C2, respectively. Between 8% and 37% of hsPRS-v2 pairs were detected by the 10 versions of GPCA, MAPPIT, KISS and Y2H when none of the hsRRS-v2 pairs were scored positive (**Fig. 2c**), which represented an average PRS detection rate of 22.8% ± 8.8% (S.D.) (**Fig. 2c**, shaded blue area).

When we considered all protein pairs identified by the union of these 10 versions corresponding to four assays, the hsPRS-v2 recovery rate reached 63% (**Fig. 2d** and **Supplementary Fig. 2d**), similar to what was obtained by combining ten distinct assays, performed in only one or two version(s), as published in the literature (65%; **Fig. 2b**). This result demonstrates that the combination of four distinct assays exploring different experimental conditions can recover as many PPIs as 10 fundamentally different techniques limited to a few versions.

### Towards developing a highly versatile binary PPI assay

In an attempt to further minimize the overall number of multi-version assays required to maximize binary PPI detection and, by extension, reduce the associated cloning efforts and lab resources, we decided to develop a highly versatile binary PPI detection system opening access to all assay versions defined by the three experimental parameters presented in **Fig. 1b**.

Although after re-analyzing published data 1.3-fold and 1.4-fold increases in PRS detection were observed when switching from one orientation to the other (*i.e.* Tag1-X/Tag2-Y to Tag1-Y/Tag2-X), or from one configuration to another (*i.e.* N1N2 to C1N2), respectively (**Supplementary Fig. 3a,b**), the impact of permuting expression systems remained unclear as no assay had been used to systematically explore this parameter. However, several high quality PPI datasets have been obtained using heterologous expression of protein pairs in yeast^12, 35^, in mammalian cells^36, 37^ and via cell-free coupled transcription-translation systems *in vitro*^38, 39^.

After surveying existing binary interaction assays, it appeared that split-protein reconstitution was the most suitable technology to cover the three experimental parameters presented in **Fig. 1b**. Indeed, compared to other binary PPI methods involving cell-specific responses or antibody-based captures, split protein-based assays enable detection of PPIs through the simple reassembly of a functional reporter protein (**Fig. 1c**). Among the different reporters available today, NanoLuc, a highly-stable 19kDa protein derived from the luciferase complex of the deep-sea shrimp *O. gracilirostris*, is gaining increasing attention for the development of binary PPI detection assays, as highlighted by recent publications^31, 32, 40, 41^. Following the single-step addition of imidazopyrazinone substrates, NanoLuc instantly produces an intense and sustainable glow-type luminescence with a specific activity about 100-fold greater than that of other luciferases, allowing its detection *in vitro* as well as in living cells and animals^31, 42, 43^. Therefore, split-NanoLuc-based technology appears to be the ideal candidate for the high-throughput detection of binary PPIs in various experimental conditions.

Among the recent studies reporting a split-NanoLuc binary PPI assay, only one extensively fragmented the enzyme to test different tag pairs in an unbiased way (**Supplementary Table 3**)^32^. After selecting a cleavage site between amino acids 156 and 157, the authors heavily modified the corresponding fragments to reduce their association affinity. The resulting moieties, 11S (156 amino acids) and 114 (11 amino acids), were then used to develop the NanoLuc Binary Technology (NanoBiT) assay^32^.

Even though NanoBiT was originally designed for expression of protein pairs in mammalian cells, we decided to evaluate whether fragments 11S and 114 could be used to further develop our new, highly versatile binary PPI detection system (**Fig. 1c**, right panel). To investigate this, we first adapted the available NanoBiT vectors with the reported original split site^32^ to achieve different tagging configurations with Gateway cloning compatibility and then benchmarked all four pairing combinations, *i.e.* N1N2, C1N2, N1C2 and C1C2, against hsPRS-v2 and hsRRS-v2. The benchmarking of these four NanoBiT versions in conditions where none of the hsRRS-v2 pairs are scored positive showed that between 3% and 25% of hsPRS-v2 pairs could be recovered (**Fig. 2c**). These results revealed optimal performances when the larger NanoBiT fragment, 11S (156 amino acids), was fused to the C-terminal extremity of the tested proteins, as demonstrated by PRS recovery rates of 22% and 25% for the C1N2 and C1C2 configurations, respectively, similar to those from four other well-established assays (**Fig. 2c**). These observations were in agreement with the author’s original assumption that the C-terminus tagging of proteins with 11S might impose minimal steric constraints^32^.

### Nanoluc 2-Hybrid (N2H): a highly versatile binary PPI assay

Although NanoBiT appears to be a powerful tool to study PPI kinetics in live cells, the comprehensive benchmarking of this assay revealed two underperforming versions, N1N2 and N1C2, that might be associated to the inherent properties of the selected tag pair^32^. Therefore, we decided to empirically define a new split site for NanoLuc in an unbiased way. To do this, we first extensively fragmented the full-length enzyme by transposon-based insertional mutagenesis (**Supplementary Fig. 4a**). Complementation of the resulting fragments, F1 and F2 (58 pairs in total), was then systematically tested using a proximity assay *in vitro* (**Supplementary Fig. 4b**), and a cleavage site between amino acids 65 and 66 was selected (**Fig. 3a**) to build the NanoLuc 2-hybrid (N2H) assay (**Fig. 1c**, right panel and **Supplementary Fig. 4c**).

**Figure 3.**
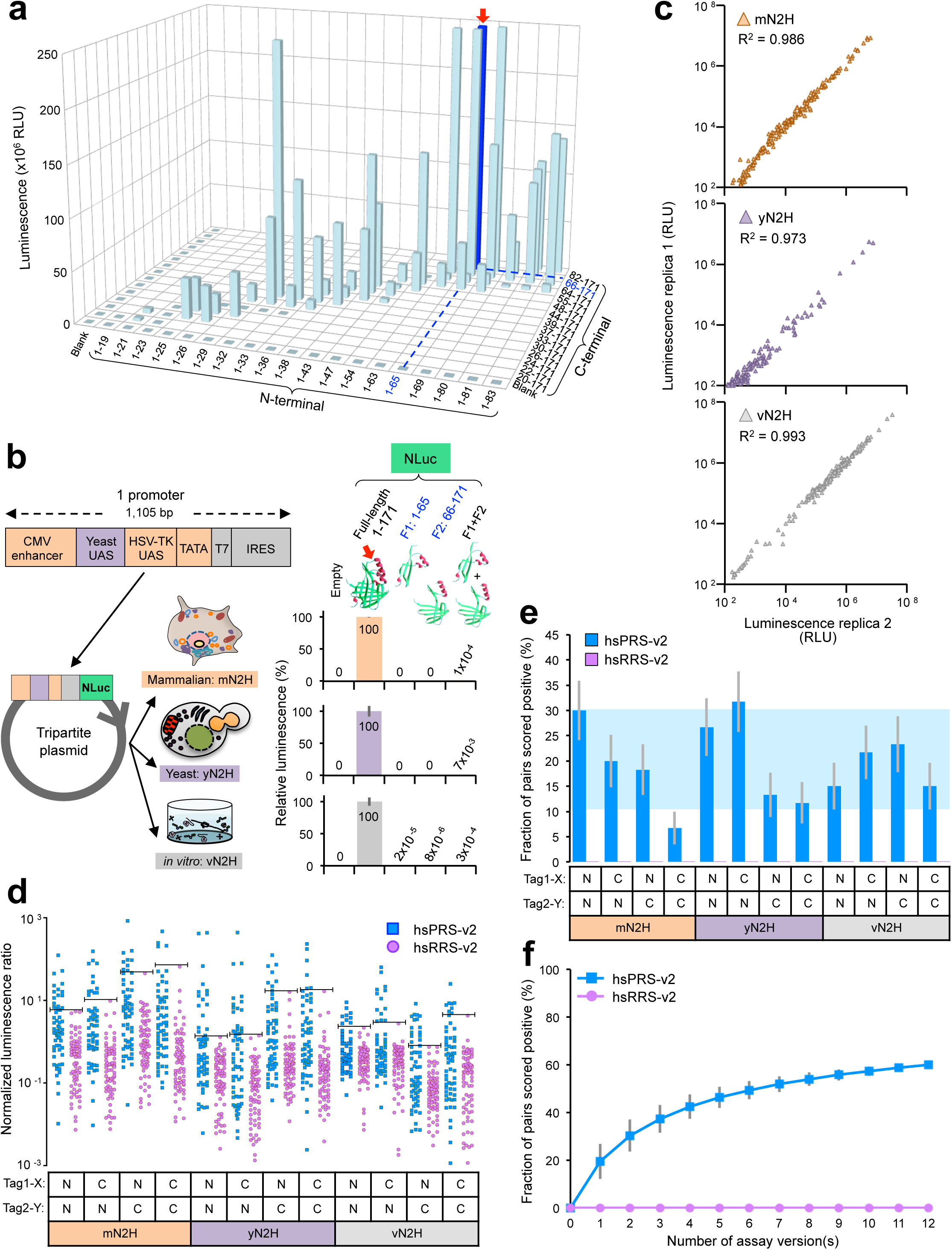
Development of the highly versatile N2H assay. (**a**) Determination of a new split site for NanoLuc. Dark blue bar and red arrow indicate the selected NanoLuc fragments. RLU=relative luminescence units. NLuc=NanoLuc. Blank control corresponds to reagents without any NanoLuc fragment. (**b**) Tripartite promoter and specific activities of selected N2H fragments, F1 and F2, normalized to full-length NanoLuc when expressed in mammalian cells (mN2H; orange), in yeast (yN2H; purple), or *in vitro* (vN2H; gray), from the same plasmid. For “empty”: no protein or fragment was expressed. CMV: cytomegalovirus; HSV-TK: herpes simplex virus-thymidine kinase; UAS: upstream activating sequence; IRES: internal ribosome entry site. Red arrow displayed on NanoLuc structure (PDB: 5IBO) indicates the cleavage area. (**c**) Reproducibility of N2H assay in three expression environments (mN2H, yN2H and vN2H) for the N1N2 tagging configuration. Each triangle in the different graphs represents a single tested protein pair. (**d**) Normalized luminescence signals for 12 versions of N2H benchmarked against hsPRS-v2 and hsRRS-v2. Blue squares and purple circles indicate PRS and RRS pairs, respectively. The threshold of no RRS pair scoring positive is indicated (horizontal line) for each assay version. (**e**) Fraction of hsPRS-v2 pairs recovered by 12 N2H assay versions when none of the hsRRS-v2 pairs are scored positive, corresponding to the thresholds displayed in (**d**). Horizontal shaded blue area indicates the average recovery rate ± S.D. for all assay versions reported in **Fig. 2c**. (**f**) Cumulative hsPRS-v2 recovery rates when versions of N2H presented in (**e**) are combined (individual data points are displayed in **Supplementary Fig. 5a**). Error bars indicate standard deviations (**b**,**f**) or standard errors of proportions (**e**).

In order to comprehensively cover the experimental parameters presented in **Fig. 1b** with the single N2H assay, we constructed four Gateway-compatible plasmids from which N2H fragments, F1 (amino acids 1-65) and F2 (amino acids 66-171), can be expressed as either N-or C-terminal fusions of any pair of proteins X and Y in yeast, in mammalian cells or *in vitro* (cell-free). To drive expression in all three environments with each individual plasmid, we engineered a tripartite promoter of ∼1kb that can easily be introduced into any vector of interest (**Fig. 3b**). The performance of this 3-in-1 promoter was evaluated by measuring the expression of full-length and fragmented NanoLuc in yeast (yN2H), in mammalian cells (mN2H) and *in vitro* (vN2H). We obtained intense signals ranging between 10^8^ and 10^9^ relative luminescence units (RLU) when the full-length NanoLuc was expressed in each individual environment. Furthermore, compared to the full-length enzyme, individually expressed NanoLuc fragments did not produce any significant signal. More importantly, the two fragments expressed together did not generate any strong self-assembly luminescence that could interfere with the accurate measurement of binary PPIs. Taken together, these results demonstrate the full functionality of both the tripartite promoter and the selected pair of NanoLuc fragments (**Fig. 3b**).

By combining NanoLuc fragments with the tripartite promoter and Gateway-cloning technology into four uniform expression vectors (pDEST-N2H-N1, -N2, -C1 and -C2), all 24 assay versions (4×3×2) presented in **Fig. 1b** could thus be implemented with the single N2H assay. The Gateway-compatibility of N2H destination plasmids also renders the entire human ORFeome collection of over 18,000 entry clones^44^ directly amenable to this assay.

As N2H was the first binary PPI detection tool in which proteins could be expressed as either N-or C-terminal fusions of protein tags in yeast (yN2H), in mammalian cells (mN2H) or *in vitro* (vN2H) from the same Gateway-compatible destination vector (**Figs. 1b,c**, **3b**), it was important to benchmark the robustness of PPI-mediated readouts in each of the three environments. Using hsPRS-v2 and hsRRS-v2 pairs, we compared bioluminescence signals between two independently replicated experiments in yN2H, mN2H and vN2H. As highlighted by the R^2^>0.97, a robust reproducibility was observed in each case when all 138 pairs from hsPRS-v2 and hsRRS-v2 were tested. In addition, a large dynamic range of luminescence readout between 10^2^ and 10^7^ RLU was consistently observed in each environment (**Fig. 3c**).

To compare the performance of N2H assay versions in detecting reference binary PPIs over random protein pairs, we normalized luminescence signals and separated hsPRS-v2 from hsRRS-v2. No major differences were observed in the overall luminescence ratios as a fraction of PRS pairs always generated higher signals than that from random pairs in each case (**Fig. 3d**). As observed for C1C2-mN2H or C1C2-yN2H, for example, the only differences in PRS detection rates between assay versions came from one or a few RRS pairs that generated higher luminescence signals than the average. These version-specific artifacts ultimately reduced the fraction of PRS pairs recovered as their corresponding thresholds of no RRS pair scoring positive increased (**Fig. 3d**). The same observation was also made for well-established assays (**Fig. 2a,c**). Thus, in conditions where none of the RRS pairs are scored positive (threshold displayed for each individual version in **Fig. 3d**), the 12 tested N2H assay versions detected between 7% and 32% of hsPRS-v2 pairs (**Fig. 3e**), demonstrating similar performances compared to five other PPI detection methods (**Fig. 2c**) that averaged 20.4% ± 9.9% (S.D.) PRS recovery (**Fig. 3e**, shaded blue area). When these 12 N2H versions were combined, a PRS detection of 60% was yielded (**Fig. 3f** and **Supplementary Fig. 5a**), similar to the 65% obtained by combining either 10 different, version-limited assays from the literature^23–28^ (**Fig. 2a,b**) or five non-N2H, multi-version assays (**Fig. 2c** and **Supplementary Fig. 5b**).

These results highlight the advantages of using a single, highly versatile PPI detection tool, such as N2H, to reduce the overall cloning efforts and lab resources (**Fig. 4a-c**), as the same set of expression vectors are used to explore several assay versions at a time (**Fig. 4d**) whilst recovering as many PPIs as a combination of many different techniques (**Fig. 4e**). The published assays in **Fig. 4** are presented in **Fig. 2a** and **Supplementary Table 1**.

**Figure 4.**
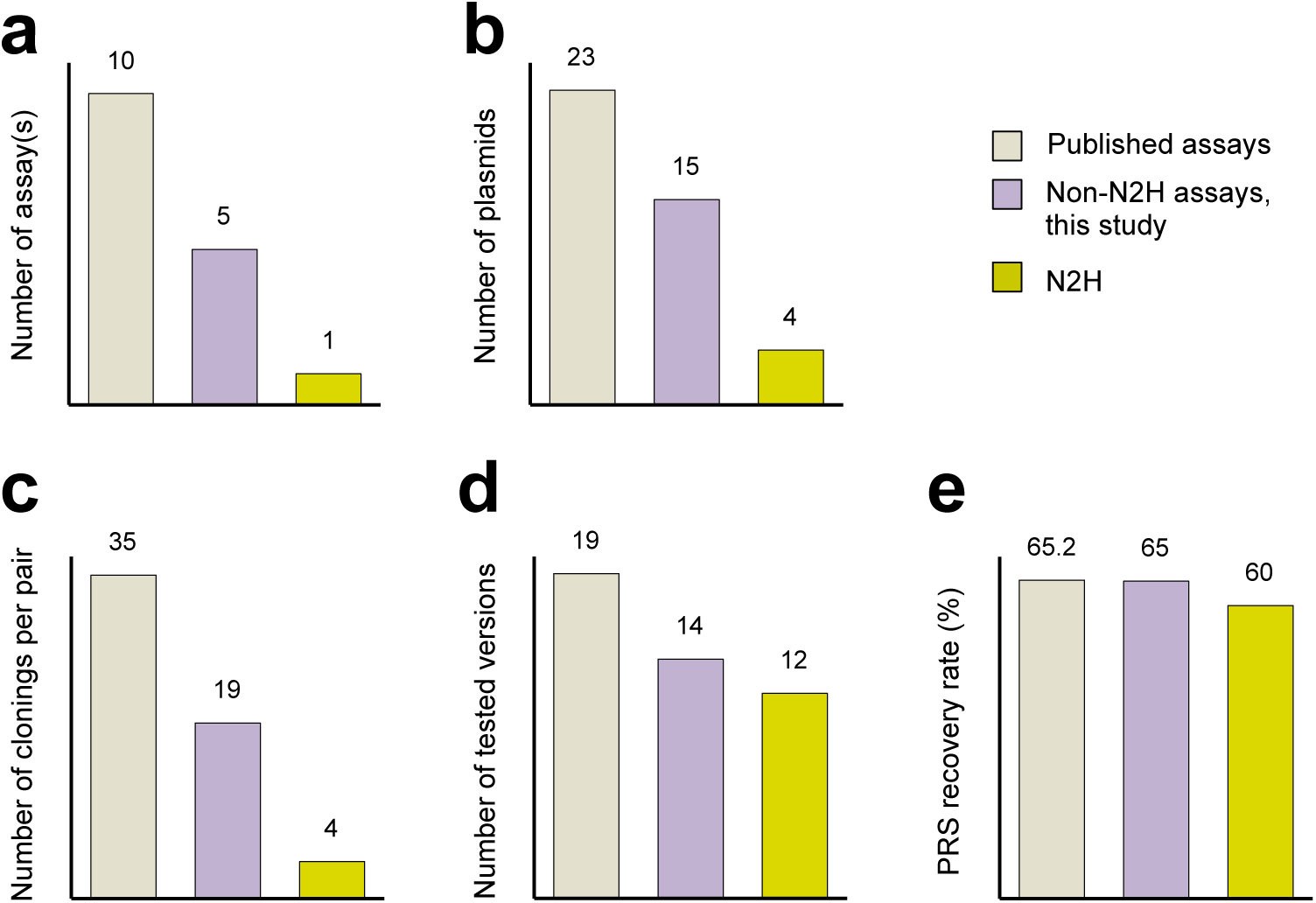
Invested resources for various combinations of binary PPI assays and PRS detection rates. Numbers of individual components required to achieve maximal PRS detections are indicated for different combinations of assays and assay versions. PRS recovery rates presented in **Fig. 4e** correspond to those found in **Fig. 2b** (published), **Supplementary Fig. 5b** (non-N2H) and **Fig. 3f** (N2H). Except for Y2H-N1C2, Y2H-C1C2 and Y2H-C1N2 for which unambiguous titrations could not be obtained from the original study^28^, all published assay versions benchmarked against hsPRS-v1 and hsRRS-v1 were used (**Fig. 2a**, **Supplementary Table 1**).

### Combining multi-version assays to maximize PPI detection

With all assays in hand to systematically evaluate the impact of the three variables presented in **Fig. 1b**, we decided to combine the different PPI assay versions previously tested against the improved reference sets (**Figs. 2c, 3e**) in respect of each individual parameter. All analyses were conducted in conditions where none of the random pairs are scored positive.

As the impact of permuting proteins X and Y from one tag to the other (Tag1-X:Tag2-Y versus Tag1-Y:Tag2-X) had already been comprehensively investigated by six different assays in the literature^23, 26, 27^ (**Supplementary Table 1**), we only repeated Y2H and MAPPIT to explore this parameter. While a single, randomly picked orientation yielded PRS recovery rates of 19% and 32%, switching to the second orientation increased detection to 25% and 40% for Y2H and MAPPIT, respectively. Importantly, cumulative PPI detection increased from 38% to 47% when a single or two orientations were combined for both assays, respectively (**Fig. 5a**). This 1.2-fold (47/38) increase was consistent with our analysis of published results^23, 26, 27^ (**Supplementary Fig. 3a**) and confirmed that permuting the orientation of tested proteins is a valuable option to increase PPI detection.

**Figure 5.**
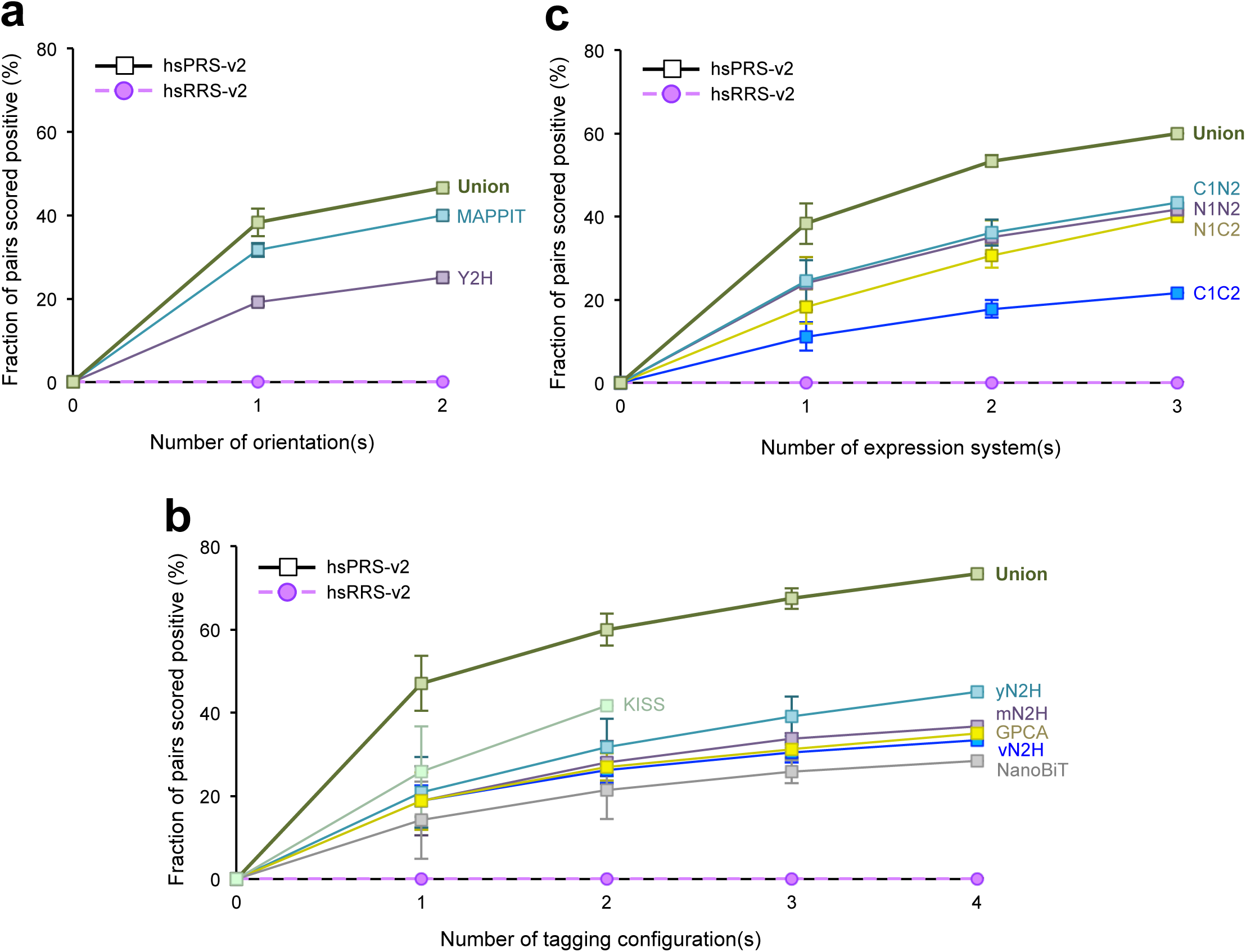
Impact of permuting assay parameters on binary PPI detection. (**a**), (**b**), (**c**) PPIs detected by permuting (**a**) protein orientations in Y2H and MAPPIT, (**b**) tagging configurations in KISS (only two versions tested), NanoBiT, GPCA, mN2H, yN2H and vN2H, and (**c**) protein expression environments in N2H. Error bars in (**a**-**c**) indicate standard deviations.

To probe the influence of different tagging configurations on PPI detection, *i.e.* the N-or C-terminal positioning of the assay tags, we systematically evaluated the contribution of each pairing combination (N1N2, C1N2, N1C2 and C1C2) with KISS, NanoBiT, GPCA, mN2H, yN2H and vN2H. For KISS, the analysis was restricted to the two published configurations: N1N2 and C1N2^25^. Using mN2H as a first example, we observed an average PRS detection rate of 19% for a single, randomly picked tagging configuration (**Fig. 5b**). When two of the four possible tagging configurations were used, the PRS recovery rate was further increased to 28%. Finally, when three and all four tagging configurations were combined, 34% and 37% of hsPRS-v2 pairs were identified (**Fig. 5b**). Thus, using more than one tagging configuration substantially increased the detection of binary PPIs by mN2H. Similar observations were made for the other assays as the PRS recovery rates were increased from an average 26% to 42% for KISS, where only two configurations were tested, and from 14% to 28% for NanoBiT, 19% to 35% for GPCA, 21% to 45% for yN2H and 19% to 33% for vN2H when only one or all four configurations were tested, respectively. The same trend was observed when combining all tested versions and a 1.6-fold increase in PRS detection obtained by extending assays from a single to multiple configurations (**Fig. 5b**, union). These results illustrate the gain of sensitivity achieved when performing the same assay with different combinations of tagging configurations, and confirm the preliminary observations made after re-analyzing published data^23, 25^ (**Supplementary Fig. 3b**).

To evaluate the impact of different protein expression systems on binary PPI detection, we implemented different versions of the N2H assay in yeast, in mammalian cells and *in vitro* (cell-free). All four tagging configurations (**Fig. 1b,c**) were systematically tested in each environment. For the N1N2 version, an average PRS detection rate of 24% was reached when performing the assay in a single environment. This recovery was increased to 35% when switching to a second expression environment. Finally, after combining detected PRS pairs from all three environments, the cumulative recovery rate reached 42% for the N1N2 versions of N2H (**Fig. 5c**). When we expanded the analysis to the other tagging configurations, N1C2, C1C2, and C1N2, we consistently observed that performing the assay with different expression systems allowed the identification of a larger fraction of PPIs (**Fig. 5c**). Indeed, a robust growth of cumulative PRS detection was observed for N1C2 (from 18% to 40%), C1C2 (from 11% to 22%) and C1N2 (from 24% to 43%) when PPIs were tested in just one or all three different expression environments, respectively. Importantly, the cumulative PPI recovery rate for all 12 N2H assay versions tested here was increased from 38% to 60% when using one or all three expression systems, respectively, which represented an overall 1.6-fold increase in PPI detection (**Fig. 5c**, union). This result also highlights that the performance of the single multi-version N2H assay (60% PRS recovery) was significantly higher than that of any other method presented in this study (**Fig. 2a,c**).

Taken together, these observations demonstrate the benefit of permuting experimental conditions in binary PPI assays to increase detection (**Fig. 5a-c**). What remained unclear was the overall complementarity of assay versions when they are all combined. To visualize the different subsets of interactions recovered by the six different assays analyzed in this study (**Figs. 2c, 3e**), we displayed the performance of their corresponding 26 assay versions in the form of a checkerboard plot. Under conditions where none of the hsRRS-v2 pairs are scored positive, the different versions detected distinct but partially overlapping subsets of PPIs, the union of which reached 78% (**Fig. 6**). Importantly, as already observed in **Fig. 3f**, the combination of 12 N2H versions detected 60% of PRS pairs, which means that this single assay recovered over three-quarters of all PPIs detected in this study (**Fig. 6**). Taken together, these results showed that a single assay exploring multiple dimensions of the parameter space (**Fig. 1b,c**) will be sufficient to map a large fraction of binary PPIs in a defined search space.

**Figure 6.**
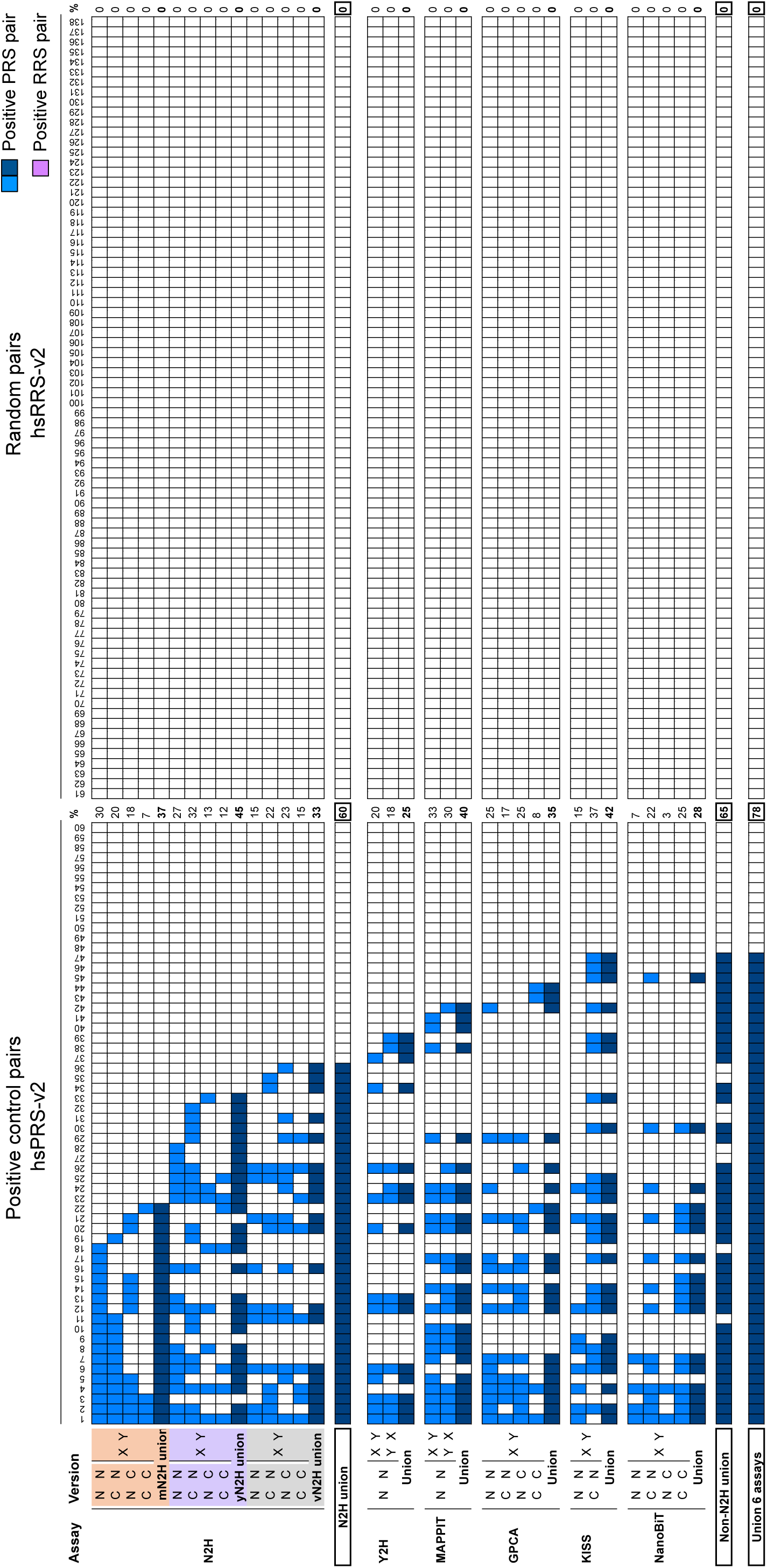
Complementarity of different assay versions. Detection of individual hsPRS-v2 pairs (left panel) by the assay versions used in **Fig. 5a-c**, under conditions where none of the hsRRS-v2 pairs (right panel) are scored positive. Detected hsPRS-v2 pairs are indicated by blue squares: light blue corresponds to individual assay versions and dark blue to the union of assay versions into distinct assays or combinations of assays. Identity of pairs can be found in **Supplementary Table 4**.

## DISCUSSION

Over the past ten years, the community has continuously released new binary PPI detection tools. The benchmarking of these assays, using a first generation of positive (hsPRS-v1) and random (hsRRS-v1) reference sets, revealed that they often identify distinct subsets of interactions^23^. Therefore, a combination of multiple techniques will be required to increase overall PPI detection and reach completeness (**Fig. 1a**). Under conditions of optimal specificity, we have shown that no single PPI assay is exceptionally superior to any other, including the most recently developed technologies. Furthermore, we have established that instead of continuously developing fundamentally different techniques, it should be possible to reach completeness by assembling a minimal number of binary PPI assays, as long as different experimental conditions, or assay versions, can be implemented for each selected assay (**Figs. 1b,c**).

We eliminated the possibility that the failure of 10 combined assays to detect about a third of known interactions in the original hsPRSv1 came from ORF clone quality issues or problematic interaction annotations by benchmarking assays against an improved, unbiased^22^ second-generation of reference sets: hsPRS-v2 and hsRRS-v2. Although the improvement of reference sets was purely focused on pair quality in this study, the next generation of PRS and RRS should include larger fractions of interactions such as those involved in higher order complex structures as well as interactions from co-complex crystal structures where direct contacts are well defined. Protein pairs for which a variety of K_D_ measurements are available^27^ will also be included to characterize assays and assay versions according to their capabilities of recovering interactions of different strengths.

Using the improved standards under conditions that maximized specificity, we have shown that considering three experimental parameters, *i.e.* protein orientation, tagging configuration and expression system (**Fig. 1b**), for assay development is a valuable option to increase PPI detection. In particular, we demonstrated that combining a relatively small number of assays exploring these parameters could lead to a PPI detection rate of up to 78%. In this context, we showed that combining 12 different versions of N2H, a new, highly versatile split NanoLuc-based assay, could recover 60% of all positive control interactions, similar to the 65% recovery rates obtained by combining either 10 published version-limited assays or four distinct multi-version assays. This means that the highly versatile N2H assay identified 77% of all PPIs detected in this study. In short, our results highlight the advantage of N2H or similar versatile systems, capable of handling several experimental parameters in a single, multi-version toolkit, to improve PPI detection and ultimately reduce the number of binary PPI assays required. This also suggests that only a few versatile binary PPI assays will be sufficient to reach complete coverage of any defined search space.

Surprisingly, 22% of hsPRS-v2 pairs remained undetected after combining the six assays tested in this study, even though all interactions had similar numbers of experimental evidence and publications (**Supplementary Fig. 6**). We examined whether there was any correlation between a tested PPI assay in this study, and the original detection method^54^ of the PRS pairs, but we did not find any correlation between these two elements. In two cases, the ORFs we used corresponded to shorter isoforms that did not contain reported interaction domains (**Supplementary Table 5**). Upon removal of these two ORFs, the PRS recovery rate by the union of all 26 assay versions yields 81%. However, two thirds of undetected hsPRS-v2 pairs contained reported interaction domains. Therefore, to reach completeness, we suggest that future assay development should be focused on expanding the number of versions covered by already-existing techniques. In that regard, we have extended our finding to the recently developed LuTHy assay^27^ and validated our original conclusions as additional PRS detection was observed when running all 16 versions (2 environments x 2 orientations x 4 configurations) allowed by this technology (data not shown). In addition, exploring yet untouched experimental parameters such as additional modifications of tag-protein fusions, or alternative expression environments including sub-cellular localization should also help reaching full coverage.

Given the difficulties of performing heterogeneous PPI assays with the necessity of preparing different clones for expression vectors and setting up different experiments, performing an equal number of N2H assay versions to acquire comparable PPI detection offers a valuable alternative to reduce these efforts (**Fig. 4a-e**).

The fact that expression environment has a strong impact on PPI detection is particularly interesting as it can easily be tuned with N2H or similar versatile assays. For example, protein pairs could be expressed in multiple cell lines, including distinct cell types and tissue origins, while a range of pH or salt concentrations could easily be tested when performing the assay *in vitro*. From a single set of plasmids, this would open access to almost unlimited assay versions to generate near-complete, high-quality binary interactome maps in a more time-effective manner.

## METHODS

### Processing publicly available hsPRS-v1 and hsRRS-v1 data

Publicly available protein interaction datasets used in this study came from the original publications^23, 24, 26^ or were provided by the authors upon request^25, 27^. Analyses were restricted to datasets for which a straightforward titration at a threshold of no RRS pair scoring positive could be applied. To evaluate the recovery rates of hsPRS-v1 when none of the RRS pairs are scored positive, data were calibrated such that the provided score for a given PRS pair must be strictly higher than the highest score of any of the RRS pairs. When authors clearly identified a particular pair as not interacting (negative) in their experiments, this assignment was also considered to generate the final dataset at a threshold where no RRS pair was scored positive. The same criterion was applied to evaluate the fractions of pairs found with available assay permutations, by using the highest RRS score found in each independent version of the corresponding assay.

### Construction of hsPRS-v2 and hsRRS-v2 reference sets

Second-generation hsPRS-v2 (60 pairs) and hsRRS-v2 (78 pairs) were constructed from pre-existing hsPRS-v1 (92 pairs) and hsRRS-v1 (92 pairs)^23^. To update the v1 sets generated over 10 years ago, we first decided to obtain single colony isolates for each ORF clone, removing all possible mutations that could have appeared in the original mini-pool ORFeome collection^11, 45^. Each entry clone passed through multiple rounds of individual single colony isolations, plasmid DNA purifications, and Sanger DNA sequencings, and only full-length, fully sequence-verified clones (next generation sequencing) were selected. By comparing the ORF sequences with genome annotations, we removed ORFs that did not match annotations in Gencode 27^46^, resulting in 76 PRS pairs and 78 RRS pairs (91.8% success rate at the ORF level, **Supplementary Fig. 2a**). We further inspected the remaining 76 PRS pairs by literature mining and informatics filtering. We revisited the supporting evidence for each PRS pair using an updated classification of binary PPI detection methods (**Supplementary Data 1**). For example, pull down and co-immunoprecipitation are no longer considered as binary PPI detection methods^12^. Ten PRS pairs not satisfying binary classification were removed. Three PRS pairs containing HLA proteins were also discarded, as no reference ORF could be defined. Finally, three PRS pairs between precursor ligands (FGF1, CXCL1, and TNFSF10) and their respective receptors were also removed due to the requirement of proteolytic cleavages of the ligands for maturation (**Supplementary Table 2** and **Supplementary Fig. 2b**). After applying these filters, at total of 60 PRS pairs and 78 RRS pairs were thus assembled into hsPRS-v2 and hsRRS-v2, respectively (**Supplementary Table 4**). Discarded pairs from hsPRS-v1 and hsRRS-v1 are presented in **Supplementary Data 1**, and PubMed IDs supporting remaining hsPRS-v2 pairs are presented in **Supplementary Data 2**. The performances of the updated reference sets were compared to the older sets by repeating four assays previously benchmarked with hsPRS-v1 and hsRRS-v1 (**Supplementary Fig. 2c**).

### Cloning ORFs into Gateway-compatible expression plasmids

Human hsPRS-v2 and hsRRS-v2 ORFs are available in pDONR221 or pDONR223 vectors, which make them compatible with the Gateway cloning technology. Each ORF was introduced into the different assay-specific expression vectors used in this study via an LR clonase-mediated Gateway reaction (Life Technologies). LR reaction products were subsequently transformed into DH5α competent bacterial cells and grown for 24 hours in ampicillin-containing TFB liquid medium. Plasmid DNA was extracted using a NucleoSpin 96 Plasmid kit from Macherey-Nagel. After PCR-amplification using plasmid-specific primers, the size of each DNA amplicon was examined by agarose gel electrophoresis. For each batch of ready-to-go destination vectors, DNA sequencing was performed on a subset of the cloned samples to check the quality of cloning before running the PPI assays.

### Construction of the tripartite promoter and N2H plasmids

The design of the 3-in-1 hybrid promoter was based on the recent description of highly potent, minimal synthetic promoters in yeast *S. cerevisiae*^47^. These promoters comprise a short synthetic upstream activating sequence (UAS) and generic core elements, including a TATA box and a Transcription Start Site (TSS). The average distance separating the UAS motif upstream of the TATA box in native yeast and mammalian cell promoters was sufficient to insert the GC boxes and the CCAAT box-binding transcription factor (CTF) binding site of the herpes simplex virus-thymidine kinase (HSV-TK) promoter^48^ between these two elements. To boost expression levels in mammalian cells, the cytomegalovirus (CMV) immediate early enhancer was inserted upstream of the yeast synthetic UAS. To drive expression *in vitro* using a coupled transcription/translation system, a T7 promoter was inserted 30 nucleotides downstream of the TATA box, immediately followed by a cricket paralysis virus internal ribosome entry site (CrPV-IRES), a versatile IRES used to initiate translation in both, mammalian and yeast cells^49, 50^. All these regulatory elements were assembled using gene synthesis (GenScript) and 40 nucleotides of sequence homology to the insertion sites in the vector backbone (pESC-LEU, Agilent, cat #217452; pESC-TRP, Agilent, cat #217453) were added for gap repair recombination in yeast^51, 52^. NanoLuc fragments, F1 (1-65) and F2 (66-171), were linked to either the N-terminal (pDEST-N2H-N1, -N2) or C-terminal (pDEST-N2H-C1,-C2) extremity of the tested protein, using a flexible hinge encoding a polypeptide of 20 amino acids along with Gateway recombinational cloning sites. Finally, expression levels in the different systems were assessed using full length or fragmented NanoLuc (Promega, cat #N1001) reporter genes (**Fig. 3b**).

### Systematic evaluation of NanoLuc fragments complementation

NanoLuc fragments, were systematically generated by transposon-based insertional mutagenesis randomly distributed between nucleotides 55 and 249 of the NanoLuc coding sequence, corresponding to amino acid residues 19 and 83, respectively (**Supplementary Fig. 4a**). Resulting moieties were characterized by Sanger DNA sequencing using pMOD forward and reverse primers according to the manufacturer’s protocol (EZ-Tn5, epicentre-Illumina, #TNP10622). Each fragment was then subcloned into a specifically designed SNAP-Tag *in vitro* expression vector, using Gateway recombination cloning (Life Technologies). They were expressed using a human IVT *in vitro* system and their complementation was assessed using a proximity-based assay.

### Design of the SNAP-Tag *in vitro* expression vector

DNA encoding the SNAP-Tag was PCR-amplified from the pSNAPf vector (NEBs, #N9183S) to be inserted in frame into the pT7CFE1-CHis plasmid provided in the human IVT kit (ThermoFisher, #88882). The Gateway cloning cassette was introduced immediately after the SNAP tag sequence using the Gateway vector conversion system (Invitrogen, # 11828-029). All subsequent cloning steps were performed using Gateway BP and LR reactions according to the manufacturer’s protocol. DNA constructs were prepared using standard maxi- and mini-prep protocols.

### Human IVT *in vitro* expression system

To assess complementation of NanoLuc fragments, a HeLa cell lysate was used for *in vitro*-coupled transcription and translation (IVT system, ThermoFisher, #88882). All the reactions were performed according to the manufacturer’s protocol, except for DNA concentrations (150 ng for both expression vectors were used) and the use of a 5x dilution of the HeLa cell lysate in the DB buffer. Concentrated DB buffer (10x) consisted in 540 mM HEPES-KOH pH: 7.6; 800 mM K-acetate; 40 mM Mg-acetate; 50 mM DTT; 5 mM Spermidine; 12.5 mM ATP and 200 mM Creatine phosphate. The final mixture containing the diluted HeLa cell lysate with the accessory proteins and the manufacturer’s reaction buffer was supplemented with 0.8 mM UTP, CTP; 1.2 mM GTP, and 50 μM of each amino acid (Promega, #L446A). T7 RNA polymerase (NEBs, #MO251L) were added at a final concentration of 1 unit/μL. Reaction was performed for 90 minutes at 30°C, directly in 96 or 384-well plates.

### *In vitro* proximity assay based on SNAP-tag capture

NanoLuc fragments fused to SNAP-tags were expressed in the human IVT system and assayed for complementation by generating a Biotin-Streptavidin complex. After IVT expression, the reaction mixture was incubated for 30 min at 30°C with 1 μL/well of a 250 μM SNAP-biotin substrate dilution in fresh DMSO. Streptavidin-biotin complexes were further generated by adding 1 μL/well of 1 mg/mL of streptavidin solution (Sigma-Aldrich, #85878). The reconstituted luciferase enzymatic activity was monitored by injecting 50 μL/well of the Nano-Glo reagent (Promega, #N1120).

### Mammalian cell-based version of N2H assay (mN2H)

HEK293T cells were seeded at 6×10^4^ cells per well in 96-well, flat-bottom, cell culture microplates (Greiner Bio-One, #655083), and cultured in Dulbecco’s modified Eagle’s medium (DMEM) supplemented with 10% fetal calf serum at 37°C and 5% CO_2_. 24 hours later, cells were transfected with 100 ng of each N2H plasmid (pDEST-N2H-N1, -N2, - C1 or -C2) using linear polyethylenimine (PEI)^46^ to co-express the protein pairs fused with complementary NanoLuc fragments, F1 and F2. The DNA/PEI ratio used for transfection was 1:3 (mass:mass). 24 hours after transfection, the culture medium was removed and 50 μL of 100x diluted NanoLuc substrate (Promega, #N1110) was added to each well of a 96-well microplate containing the transfected cells. Plates were incubated for 3 minutes at room temperature. Luciferase enzymatic activity was measured using a TriStar or CentroXS luminometer (Berthold; 2 seconds integration time).

The stock solution of PEI HCl (PEI MAX 40000; Polysciences Inc; Cat# 24765) was prepared according to the manufacturer’s instructions. Briefly, 200 mg of PEI powder added to 170 mL of water, stirred until complete dissolution, and pH was adjusted to 7 with 1 M NaOH. Water was added to obtain a final concentration of 1 mg/mL, and the stock solution was filtered through a 0.22 μm membrane.

### Yeast cell-based version of N2H assay (yN2H)

Using a high-efficiency LiAc/SS carrier DNA/PEG protocol^53^, haploid *S. cerevisiae* Y8800 (*MATa, leu2-3,112 trp1-901, his3Δ200, ade2-101 ura3-52, gal4Δ, gal80Δ, chy2^R^, GAL2::ADE2, GAL7::LacZ@met2, GAL1::HIS3@LYS2*) and Y8930 (*MATα, leu2-3,112 trp1-901, his3Δ200, ade2-101 ura3-52, gal4Δ, gal80Δ, chy2^R^, GAL2::ADE2, GAL7::LacZ@met2, GAL1::HIS3@LYS2*) yeast strains were transformed with pDEST-N2H plasmids (pDEST-N2H-N1, -N2, -C1 and –C2) containing a *TRP1* (N2/C2/Fragment 2 vectors) or a *LEU2* (N1/C1/Fragment 1 vectors) cassette, respectively. For each individual transformation, 0.3 to 1.5 μg of high-purity plasmid and 1 mL of log phase yeast cell culture (OD_600 nm_ ∼ 0.6-0.9) were used. The resulting transformants carrying N2/C2/Fragment2 vectors (*TRP1* cassette) or N1/C1/Fragment1 vectors (*LEU2* cassette) were spotted (8 μL per spot) onto solid synthetic complete media lacking tryptophan (SC-W) or leucine (SC-L), respectively. After 3 days of incubation at 30°C, each transformant growing on solid SC-W or SC-L was picked by scratching the corresponding spot and inoculated into a 96-well microplate (Fisher Scientific/Corning, #07200720A) containing 160 μL of liquid SC-W or SC-L per well, respectively. Inoculated yeast cells were incubated overnight (15 to 18 hours), at 30°C and under shaking conditions (220 rpm). Haploid yeast were mated by inoculating 5 μL of transformed *MATa* Y8800 and 5 μL of transformed *MATα* Y8930 into a 96-well microplate containing 160 μL of YEPD medium (yeast extract, peptone, dextrose) per well. Mating was obtained by growing cells overnight (15 to 18 hours), at 30°C and under shaking conditions (220 rpm). A first diploid selection was conducted by inoculating 10 μL of mated yeast cells into a 96-well microplate containing 160 μL of SC-LW medium per well. Cells were incubated overnight (15 to 18 hours), at 30°C and under shaking conditions (220 rpm). A second diploid selection was conducted by inoculating 50 μL from the first diploid selection culture into a 96-well deep well plate (QIAGEN, #19579) containing 1.2 mL of liquid SC-LW medium per well. Diploids were grown by incubating cells for 24 hours, at 30°C and under shaking conditions (900 rpm). Cell density was determined by transferring 100 μL of each diploid cell culture into individual wells of a 96-well microplate and measuring OD_600 nm_ with a microplate reader (optional). Deep well plates containing diploid yeast cells were centrifuged at 2,500g for 15 minutes. After centrifugation, the supernatant was discarded and residual SC-LW medium removed by gently blotting deep well plates with absorbent paper. Each yeast cell pellet was individually resuspended into 100 μL of the NanoLuc Assay solution by gently pipetting up-and-down several times, until cell pellets were fully dispersed. Homogenized solutions were transferred into white flat-bottom 96-well plates (Greiner bio-one, cat# 655073) that were then sealed with aluminum foil and incubated for one hour at room temperature. After incubation, plates were shaken for 30 seconds at 950 rpm to resuspend yeast cells. Using a luminometer (Berthold) set up for one-second orbital shaking before each measurement (speed: fast; diameter: 5 mm), luminescence was evaluated for each reaction. Integration time of the luminescence signal was set at 2 seconds per sample.

### *In vitro* version of N2H assay (vN2H)

Synthesis of proteins was performed separately using the *in vitro* coupled transcription-translation system based on Insect Cell Extract (ICE) (TnT-T7-ICE, Promega, #L1102). In this ICE coupled transcription-translation system, transcription is driven by T7 RNA-Polymerase present in the ICE reagent. The necessary T7 promoter is included in the N2H synthetic promoter as described in **Fig. 3b**. Each reaction was performed in 25 μL final volume with 1 μg of pDEST-N2H plasmid (5 μL at 200 ng/μL) and 20 μL of diluted insect cell extract (containing 4 μL of ICE, 3.2 μL of 5x ICB dilution buffer and 8 units of T7-RNAP-Promega#P4074, supplemented with H_2_O up to 20 μL). Incubation was at 30°C for 4 hours. The synthesis reaction was stopped by addition of 50 μL of 1x PBS (Gibco, #14190-094) containing 20% glycerol (for storage at −80°C). The NanoLuc assay was carried out in 96-well format by mixing 5 μL of each *in* vitro synthesized protein with 15 μL of 1x PBS per well. After incubation at 30°C for 90 minutes, 50 μL of 100x diluted NanoLuc substrate (Promega, #N1110) were added, and luciferase enzymatic activity was measured using a TriStar or CentroXS luminometer (Berthold; 5 seconds integration time).

Concentrated insect cell buffer (ICB 5x) consists of 150 mM HEPES-KOH pH: 7.6; 750 mM K-acetate; 19.5 mM Mg-acetate; 12.5 mM DTT; 500 μM spermidine; 100 mM creatine phosphate; 0.5 mg/mL creatinine kinase; 8.75 mM ATP; 1.5 mM GTP, CTP, UTP and 500 μM amino acids mix. The stock solution of amino acids (3.4 mM) was made by dissolving the 20 powders into 5 M KOH, at concentrations ranging from 1.6 to 4 M. The resulting mixtures containing each of the 20 amino acids were diluted in water to achieve a final concentration of 238 mM. The final aqueous solution was adjusted to pH 7.5 with acetic acid, diluted to 3.4 mM and frozen in liquid nitrogen for storage at −80°C.

### Yeast 2-Hybrid (Y2H) assay

Using a high-efficiency LiAc/SS carrier DNA/PEG protocol^53^, haploid *S. cerevisiae* Y8800 (*MATa, leu2-3,112 trp1-901, his3Δ200, ade2-101 ura3-52, gal4Δ, gal80Δ, chy2^R^, GAL2::ADE2, GAL7::LacZ@met2, GAL1::HIS3@LYS2*) and Y8930 (*MATα, leu2-3,112 trp1-901, his3Δ200, ade2-101 ura3-52, gal4Δ, gal80Δ, chy2^R^, GAL2::ADE2, GAL7::LacZ@met2, GAL1::HIS3@LYS2*) yeast strains were transformed with pDEST-AD and pDEST-DB plasmids^16^ containing a *TRP1* or a *LEU2* cassette, respectively. For auto-activation tests (DB-X/Y *vs.* AD-empty), Y8800 (*MATa*) yeast cells were also transformed with a pDEST-AD-empty plasmid. Six standard Y2H controls were also transformed into yeast cells: 1) pDEST-AD-empty/pDEST-DB-empty, 2) pDEST-AD-CBLB/pDEST-DB-GRB2, 3) pDEST-AD-XIAP/pDEST-DB-CASP9, 4) pDEST-AD-empty/pDEST-DB-scGal4, 5) pDEST-AD-empty/pDEST-DB-WDR62, and 6) pDEST-AD-scUme6 (a.a. 531-a.a. 836)/pDEST-DB. For each individual transformation, 0.3 to 1.5 μg of high-purity plasmid and 1 mL of a log phase-growth yeast cell culture (OD_600 nm_ ∼ 0.6-0.9) were used. The resulting transformants carrying pDEST-AD vectors (*TRP1* cassette) or pDEST-DB vectors (*LEU2* cassette) were spotted (8 μL per spot) onto solid synthetic complete media lacking tryptophan (SC-W) or leucine (SC-L), respectively. After 3 days of incubation at 30°C, each transformant growing on solid SC-W or SC-L was picked by scratching the corresponding spot, and inoculated into a 96-well costar plate (Fisher Scientific/Corning, #07200720A) containing 160 μL of liquid SC-W or SC-L per well, respectively. Inoculated yeast cells were incubated overnight (15 to 18 hours), at 30°C and under shaking conditions (220 rpm). Haploid yeast were mated by inoculating 5 μL of transformed *MATa* Y8800 and 5 μL of transformed *MATα* Y8930 into a 96-well microplate containing 160 μL of YEPD medium (yeast extract, peptone, dextrose) per well. The exact same mating was done for the auto-activation tests (DB-X/Y-containing cells mated with AD-empty-containing cells). Mating was obtained by growing cells overnight (15 to 18 hours), at 30°C and under shaking conditions (220 rpm). The diploid selection was conducted by inoculating 10 μL of mated yeast cells into a 96-well microplate containing 160 μL of SC-LW medium per well. Cells were incubated overnight (15 to 18 hours), at 30°C and under shaking conditions (220 rpm). Using a liquid handling robot, each diploid cell culture was then spotted in quadruplicate (1 μL/spot) onto two different solid media plates: 1) SC-LW, and 2) SC-LW lacking histidine and containing 1 mM of 3-Amino-1,2,4-triazole (3-AT) (SC-LWH, 1 mM 3-AT). The six controls described above were also spotted on each solid plate to validate composition of selective media (based on their known phenotypes on these media) and compare their growth rates to the ones of tested protein pairs. Plates were incubated for four days at 30°C and two days at room temperature before evaluating growth scores.

### *Gaussia princeps* complementation assay (GPCA)

HEK293T cells were cultured as described for the mN2H assay version. For each protein, 100 ng of purified plasmid DNA was transfected into HEK293T cells in 96-well, flat-bottom, cell culture plates (Greiner Bio-One, #655083) supplemented with 10%FBS in DMEM using polyethylenimine^24^. The DNA/PEI ratio (mass:mass) was 1:3. GPCA vectors carry the human cytomegalovirus (CMV) promoter and are maintained as high copy number with the human virus SV40 replication origin in mammalian cells. 24 hours after DNA transfection, the cell culture medium was removed, cells were gently washed with 150 μL pre-warmed 1x PBS, 40 μL of lysis buffer was then added per well, and cell lysis was performed under vigorous shaking of the plate for 20 minutes at 900 rpm. Luminescence was measured by auto-injecting 50 μL Renilla luciferase substrate (Renilla Luciferase Assay system, catalog No. E2820, Promega) per well and integrating light output for 4 seconds using a TriStar luminometer (Berthold).

### Construction of C-terminal plasmids for GPCA

GPCA C1 and C2 vectors are based on two fragments of the humanized *Gaussia princeps* luciferase (herein referred as GLuc), similar to the GPCA N1 and N2 vectors^24^. Both GLuc fragments were linked to the C-terminal extremity of the tested proteins by a flexible hinge polypeptide of 20 amino acids, including the Gateway recombinational cloning sites. To normalize mRNA translation initiation, a consensus Kozak translation start sequence was added. Both constructs were carried by the same CMV driven mammalian expression vector (pCI-neo derived, Promega) and were maintained as multi-copies within the cells via the presence of the SV40 replication origin.

### NanoLuc^®^ Binary Technology (NanoBiT) assay

HEK293T cells were cultured as described for the mN2H assay version. For each protein, 100 ng of purified plasmid DNA was transfected into HEK293T cells grown in 96-well, flat-bottom, cell culture microplates (Greiner Bio-One, #655083). The DNA/PEI ratio (mass:mass) was 1:3 for the transfection. 24 hours later, 100 μL of the NanoLuc substrate (Promega, #N1110) were added to each well, plates were incubated for 3 minutes at room temperature, and luminescence output was measured using a TriStar luminometer (Berthold; 1 second integration time).

### Construction of Gateway-compatible plasmids for NanoBiT

The Gateway reading frame cassette was amplified by PCR, and polished with a PCR polishing kit (Agilent, # 200409) to generate blunt-end DNA. NanoBiT vectors (Promega, # N2014) with N-and C-terminus tagging configurations (NB MCS1, NB MCS2, NB MCS3, NB MCS4) were obtained from Promega, each containing an HSV-TK promoter for protein expression in mammalian cells and an ampicillin resistance marker for bacterial selection of transformed cells. Each plasmid DNA for NanoBiT vectors (NB MCS1-4) was digested with a single-cut restriction enzyme (EcoRI for NB MCS1 and 2, and SacI for NB MCS3 and 4) at the multiple cloning sites (MCS). The linearized plasmid DNA was processed with the PCR polishing kit generating blunt-end DNA. Antarctic phosphatase (NEB, M0289S) was used to remove 5’-phosphates to reduce self-ligation background of the linearized vector. The ligation reaction was performed with T4 DNA ligase (NEB, M0202S) joining the linearized NanoBiT vectors (NB MCS1-4) with the polished Gateway reading frame cassette. The ligated, Gateway compatible NanoBiT vectors (NanoBiT-GW-N1, -N2, -C1 and –C2) were introduced into One Shot^TM^ ccdB survival^TM^ 2 T1^R^ competent *E. coli* cells (Invitrogen, #A10460) via heat shock bacterial transformation, and selected on chloramphenicol-containing LB agar plates. The sequences of constructed NanoBiT-GW vectors were examined by Sanger DNA sequencing. The final vectors contained correct sequences of the promoter, NanoBiT fragments, junctions, and Gateway cassette.

### Mammalian protein-protein interaction trap (MAPPIT) and Kinase substrate sensor (KISS) assays

HEK293T cells were cultured as described for the mN2H assay version. In both MAPPIT and KISS assays^12, 25^, HEK293T cells were transfected with bait, prey and reporter plasmids applying a standard calcium phosphate transfection method. Luciferase activity was measured 48 hours after transfection using the Luciferase Assay System kit (Promega) on a Enspire luminometer (Perkin-Elmer). For MAPPIT, half of the cells were stimulated with erythropoietin (2 ng/mL; GenScript) 24 hours after transfection. For each tested pair in MAPPIT, the fold-induction value (signal from 2 stimulated culture wells divided by the signal from 2 unstimulated culture wells) was calculated. For KISS the average of 4 culture wells was used.

### Scoring hsPRS-v2 pairs

Data for each assay version performed in this study are presented in **Source Data 1**. All binary PPI experiments were independently performed three times. For the different versions of N2H experiments, the average signal of the three replicates was used to determine a raw luminescence value for each protein pair X/Y. Control experiments were performed similarly where protein X or Y fused to the NanoLuc fragment F1 or F2 was co-expressed with the matching NanoLuc fragment alone (X/empty or empty/Y). For each protein pair X-Y, we calculated a normalized luminescence ratio (NLR) corresponding to the raw luminescence value of the tested pair (X-Y) divided by the maximum luminescence value from one of the two controls (X-empty or empty-Y), as summarized in equation (1) below^25^.

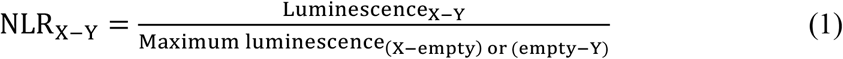

To identify protein pairs that score positive at a threshold of no RRS detection, two criteria were considered: 1) NLR_PRS_ > NLR_HIGHEST RRS_, and 2) NLR_PRS_ > 1.0. All PRS pairs that did not meet these two criteria were not scored and therefore defined as not detectable in the corresponding assay version. The same analysis and criteria were applied for both KISS and MAPPIT assays.

For GPCA and NanoBiT experiments, a PRS pair was scored positive if its corresponding raw luminescence signal (from the average of three independent experiments) was strictly higher than the RRS pair having the highest raw luminescence signal (from the average of three independent experiments).

For Y2H assay, diploid yeast cells containing both, DB-X/Y bait and AD-Y/X prey plasmids, were spotted in quadruplicate on selective medium supplemented with 3-AT (SC-LWH, 3-AT). After titration of 3-AT, a minimal concentration of 1 mM was selected as no RRS pairs were detected for this specific threshold^12^. In parallel, diploid yeast cells containing DB-X/Y bait and AD-empty prey vectors were also used to identify bait proteins that are auto-activators of the *HIS3* selection marker. After 6 days of culture, yeast growth was visually determined and scored from “0” (no growth) to “4” (strong growth) for both the DB-X/Y:AD-Y/X and DB-X/Y:AD-empty combinations. If the growth score was >0 for the DB-X/Y:AD-Y/X combination in at least two of the three experiments performed and was superior by at least 2 units to the one of DB-X/Y:AD-empty combination, a protein pair was scored positive.

### Construction of cumulative recovery plots

To plot the fraction of interaction recovered for *n* number of assays or assay versions, the recovery rate for each of the possible *n* combinations was calculated first (**Supplementary Figs. 1c, 2d, 5a**) and then averaged (**Figs. 2b,d, 3f** and **Supplementary Fig. 5b**). The error bars in **Figs. 2b,d, 3f** and **Supplementary Fig. 5b** indicate the standard deviation of the average.

## DATA AVAILABILITY

The data supporting the findings from this study are included either in the manuscript and its associated supplementary files, or can be extracted from the original referenced publications. The protein-protein interaction data have been deposited to the IntAct database (https://www.ebi.ac.uk/intact/), and will be publicly available with PubMed ID upon publication. All data are also available from the authors upon request. N2H plasmids and their corresponding maps were deposited to the Addgene plasmid repository (https://www.addgene.org/Marc_Vidal/).

## ACKNOWLEDGEMENTS

The authors would like to thank all members of the Vidal and Jacob laboratories for helpful discussions throughout this project. The authors thank Pascal Braun, and Mikko Taipale for providing inputs on the experimental procedures of various binary PPI assays, and Steffi de Rouck for technical assistance. This work was supported by a Claudia Adams Barr Award to S.G.C., a Fonds de la Recherche Scientifique (F.R.S.-FNRS)-Télévie Grant and a Wallonia-Brussels International (WBI)-World Excellence Fellowship to J.O., NHGRI grants U41HG001715 awarded to M.V., D.E.H. and M.A.C, and P50HG004233 awarded to M.V. We acknowledge the support of Labex IBEID (10-LABX-0062). M.V. is a Chercheur Qualifié Honoraire from the Fonds de la Recherche Scientifique (FRS-FNRS, Wallonia-Brussels Federation, Belgium).

## AUTHOR CONTRIBUTIONS

S.G.C., J.O., P.-O.V., M.V., and Y.J. conceived the idea. S.G.C., J.O., P.C., P.-O.V., K.S., I.L., M.D.S., C.D., S.R., E.P.C., Y.L.J., and Y.J. performed the experiments and collected the data. S.G.C., J.O., P.C., P-O.V., K.L., L.L., I.L., W.B., J.D.L.R., J.-C.T., T.H., and Y.J. analyzed the data. L.J. performed structural modeling. K.L., S.V.D.W., P.T., E.E.W., J.T., J.-C.T., T.H., D.E.H., M.V. and M.A.C. provided critical insights. S.G.C., J.O., P.-O. V., D.E.H., M.V., M.A.C., and Y.J. wrote the manuscript.

## COMPETING INTERESTS

The authors declare no competing interests.

## Notes

#### Summary of Updates

We updated the manuscript to reflect comments from the reviewers. This version was submitted for final publication.

## REFERENCES

1. A reference standard for genome biology. Nat. Biotechnol. 36, 1121 (2018).

2. Harper, J.W. & Bennett, E.J. Proteome complexity and the forces that drive proteome imbalance. Nature 537, 328–338 (2016).

3. Riley, N.M. et al. The negative mode proteome with activated ion negative electron transfer dissociation (AI-NETD). Mol. Cell. Proteomics 14, 2644–2660 (2015).

4. Chayen, N.E. & Saridakis, E. Protein crystallization: from purified protein to diffraction-quality crystal. Nat. Methods 5, 147–153 (2008).

5. Liu, H.L. & Hsu, J.P. Recent developments in structural proteomics for protein structure determination. Proteomics 5, 2056–2068 (2005).

6. Yee, A.A. et al. NMR and X-ray crystallography, complementary tools in structural proteomics of small proteins. J. Am. Chem. Soc. 127, 16512–16517 (2005).

7. Wang, H.W. & Wang, J.W. How cryo-electron microscopy and X-ray crystallography complement each other. Protein Sci. 26, 32–39 (2017).

8. Vidal, M., Cusick, M.E. & Barabási, A.L. Interactome networks and human disease. Cell 144, 986–998 (2011).

9. Luck, K., Sheynkman, G.M., Zhang, I. & Vidal, M. Proteome-scale human interactomics. Trends Biochem. Sci. 42, 342–354 (2017).

10. Stelzl, U. et al. A human protein-protein interaction network: a resource for annotating the proteome. Cell 122, 957–968 (2005).

11. Rual, J.F. et al. Towards a proteome-scale map of the human protein-protein interaction network. Nature 437, 1173–1178 (2005).

12. Rolland, T. et al. A proteome-scale map of the human interactome network. Cell 159, 1212–1226 (2014).

13. Huttlin, E.L. et al. The BioPlex network: a systematic exploration of the human interactome. Cell 162, 425–440 (2015).

14. Hein, M.Y. et al. A human interactome in three quantitative dimensions organized by stoichiometries and abundances. Cell 163, 712–723 (2015).

15. Wan, C. et al. Panorama of ancient metazoan macromolecular complexes. Nature 525, 339–344 (2015).

16. Walhout, A.J. et al. Protein interaction mapping in *C. elegans* using proteins involved in vulval development. Science 287, 116–122 (2000).

17. Tewari, M. et al. Systematic interactome mapping and genetic perturbation analysis of a *C. elegans* TGF-beta signaling network. Mol. Cell 13, 469–482 (2004).

18. Flores, A. et al. A protein-protein interaction map of yeast RNA polymerase III. Proc. Natl Acad. Sci. USA 96, 7815–7820 (1999).

19. De Las Rivas, J. & Fontanillo, C. Protein-protein interaction networks: unraveling the wiring of molecular machines within the cell. Brief Funct. Genomics 11, 489–496 (2012).

20. Cowley, M.J. et al. PINA v2.0: mining interactome modules. Nucleic Acids Res. 40, D862–865 (2012).

21. Fields, S. & Song, O. A novel genetic system to detect protein-protein interactions. Nature 340, 245–246 (1989).

22. Venkatesan, K. et al. An empirical framework for binary interactome mapping. Nat. Methods 6, 83–90 (2009).

23. Braun, P. et al. An experimentally derived confidence score for binary protein-protein interactions. Nat. Methods 6, 91–97 (2009).

24. Cassonnet, P. et al. Benchmarking a luciferase complementation assay for detecting protein complexes. Nat. Methods 8, 990–992 (2011).

25. Lievens, S. et al. Kinase substrate sensor (KISS), a mammalian in situ protein interaction sensor. Mol. Cell. Proteomics 13, 3332–3342 (2014).

26. Trepte, P. et al. DULIP: a dual luminescence-based co-immunoprecipitation assay for interactome mapping in mammalian cells. J. Mol. Biol. 427, 3375–3388 (2015).

27. Trepte, P. et al. LuTHy: a double-readout bioluminescence-based two-hybrid technology for quantitative mapping of protein-protein interactions in mammalian cells. Mol. Syst. Biol. 14, e8071 (2018).

28. Chen, Y.C., Rajagopala, S.V., Stellberger, T. & Uetz, P. Exhaustive benchmarking of the yeast two-hybrid system. Nat. Methods 7, 667–668 (2010).

29. Caufield, J.H., Sakhawalkar, N. & Uetz, P. A comparison and optimization of yeast two-hybrid systems. Methods 58, 317–324 (2012).

30. Vidal, M. & Fields, S. The yeast two-hybrid assay: still finding connections after 25 years. Nat. Methods 11, 1203–1206 (2014).

31. Hall, M.P. et al. Engineered luciferase reporter from a deep sea shrimp utilizing a novel imidazopyrazinone substrate. ACS Chem. Biol. 7, 1848–1857 (2012).

32. Dixon, A.S. et al. NanoLuc complementation reporter optimized for accurate measurement of protein interactions in cells. ACS Chem. Biol. 11, 400–408 (2016).

33. Walhout, A.J. et al. GATEWAY recombinational cloning: application to the cloning of large numbers of open reading frames or ORFeomes. Methods Enzymol. 328, 575–592 (2000).

34. Sahni, N. et al. Widespread macromolecular interaction perturbations in human genetic disorders. Cell 161, 647–660 (2015).

35. Stellberger, T. et al. Improving the yeast two-hybrid system with permutated fusions proteins: the Varicella Zoster Virus interactome. Proteome Sci. 8, 8 (2010).

36. Lievens, S. et al. Array MAPPIT: high-throughput interactome analysis in mammalian cells. J. Proteome Res. 8, 877–886 (2009).

37. Riegel, E., Heimbucher, T., Hofer, T. & Czerny, T. A sensitive, semi-quantitative mammalian two-hybrid assay. Biotechniques 62, 206–214 (2017).

38. Tang, Y., Qiu, J., Machner, M. & LaBaer, J. Discovering protein-protein interactions using nucleic acid programmable protein arrays. Curr. Protoc. Cell Biol. 74, 15.21.1.-15.21.14 (2017).

39. Yazaki, J., Galli, M., Kim, A.Y. & Ecker, J.R. Profiling interactome networks with the HaloTag-NAPPA in situ protein array. Curr. Protoc. Plant Biol. 3, e20071 (2018).

40. Verhoef, L.G., Mattioli, M., Ricci, F., Li, Y.C. & Wade, M. Multiplex detection of protein-protein interactions using a next generation luciferase reporter. Biochim. Biophys. Acta 1863, 284–292 (2016).

41. Mo, X. et al. AKT1, LKB1, and YAP1 revealed as MYC interactors with NanoLuc-based protein-fragment complementation assay. Mol. Pharmacol. 91, 339–347 (2017).

42. Stacer, A.C. et al. NanoLuc reporter for dual luciferase imaging in living animals. Mol. Imaging 12, 1–13 (2013).

43. Germain-Genevois, C., Garandeau, O. & Couillaud, F. Detection of brain tumors and systemic metastases using NanoLuc and FLuc for dual reporter imaging. Mol. Imaging Biol. 18, 62–69 (2016).

44. ORFeome Collaboration. The ORFeome Collaboration: a genome-scale human ORF-clone resource. Nat. Methods 13, 191–192 (2016).

45. Lamesch, P. et al. *C. elegans* ORFeome version 3.1: increasing the coverage of ORFeome resources with improved gene predictions. Genome Res. 14, 2064–2069 (2004).

46. Harrow, J. et al. GENCODE: the reference human genome annotation for The ENCODE Project. Genome Res. 22, 1760–1774 (2012).

47. Redden, H. & Alper, H.S. The development and characterization of synthetic minimal yeast promoters. Nat. Commun. 6, 7810 (2015).

48. Jones, K.A., Yamamoto, K.R. & Tjian, R. Two distinct transcription factors bind to the HSV thymidine kinase promoter in vitro. Cell 42, 559–572 (1985).

49. Thompson, S.R., Gulyas, K.D. & Sarnow, P. Internal initiation in *Saccharomyces cerevisiae* mediated by an initiator tRNA/eIF2-independent internal ribosome entry site element. Proc. Natl Acad. Sci. USA 98, 12972–12977 (2001).

50. Fernández, I.S., Bai, X.C., Murshudov, G., Scheres, S.H. & Ramakrishnan, V. Initiation of translation by cricket paralysis virus IRES requires its translocation in the ribosome. Cell 157, 823–831 (2014).

51. Orr-Weaver, T.L., Szostak, J.W. & Rothstein, R.J. Genetic applications of yeast transformation with linear and gapped plasmids. Methods Enzymol. 101, 228–245 (1983).

52. Walhout, A.J. & Vidal, M. High-throughput yeast two-hybrid assays for large-scale protein interaction mapping. Methods 24, 297–306 (2001).

53. Yu, H. et al. High-quality binary protein interaction map of the yeast interactome network. Science 322, 104–110 (2008).

54. Cusick, M.E. et al. Literature-curated protein interaction datasets. Nat. Methods 6, 39–46 (2009).

