## Supplementary Text and Figures for "Maximizing binary interactome mapping with a minimal number of assays"

[1]Center for Cancer Systems Biology (CCSB), Dana-Farber Cancer Institute (DFCI), 450 Brookline Avenue, Boston, MA 02215, USA. [2]Department of Genetics, Blavatnik Institute, Harvard Medical School (HMS), 77 Avenue Louis Pasteur, Boston, MA 02115, USA. [3]Department of Cancer Biology, Dana-Farber Cancer Institute, 450 Brookline Avenue, Boston, MA 02215, USA. [4]Laboratory of Viral Interactomes, Unit of Molecular Biology of Diseases, Groupe Interdisciplinaire de Génomique Appliquée (GIGA Institute), University of Liège, 7 Place du 20 Août, 4000 Liège, Belgium. [5]Département de Virologie, Unité de Génétique Moléculaire des Virus à ARN (GMVR), Institut Pasteur, UMR3569, Centre National de la Recherche Scientifique (CNRS), Université Paris Diderot, Sorbonne Paris Cité, 28 rue du Docteur Roux, 75015 Paris, France. [6]Équipe Chimie, Biologie, Modélisation et Immunologie pour la Thérapie (CBMIT), Laboratoire de Chimie et Biochimie Pharmacologiques et Toxicologiques (LCBPT), Centre Interdisciplinaire Chimie Biologie-Paris (CICB-Paris), UMR8601, CNRS, Université Paris Descartes, 45 rue des Saints-Pères, 75006 Paris, France. [7]Center for Medical Biotechnology, Vlaams Instituut voor Biotechnologie (VIB), 3 Albert Baertsoenkaai, 9000 Ghent, Belgium. [8]Cytokine Receptor Laboratory (CRL), Department of Biomolecular Medicine, Faculty of Medicine and Health Sciences, Ghent University, 3 Albert Baertsoenkaai, 9000 Ghent, Belgium. [9]Centre de Bioinformatique, Biostatistique et Biologie Intégrative (C3BI), Institut Pasteur, 28 rue du Docteur Roux, 75015 Paris, France. [10]Département de Biologie Structurale et Chimie, Unité de Chimie et Biocatalyse, Institut Pasteur, UMR3523, CNRS, 28 rue du Docteur Roux, 75015 Paris, France. [11]Neuroproteomics, Max Delbrück Center for Molecular Medicine, 10 Robert-Rössle-Str., 13125 Berlin, Germany. [12]Brain Development and Disease, Institute of Molecular Biotechnology of the Austrian Academy of Sciences (IMBA), 3 Dr. Bohr-Gasse, 1030 Vienna, Austria. [13]Cancer Research Center (CiC-IBMCC, CSIC/USAL), Consejo Superior de Investigaciones Científicas (CSIC), University of Salamanca (USAL), Campus Miguel de Unamuno, 37007 Salamanca, Spain. [14]These authors contributed equally: Soon Gang Choi, Julien Olivet, Patricia Cassonnet, Pierre-Olivier Vidalain.

⋆

### SUPPLEMENTARY FIGURES

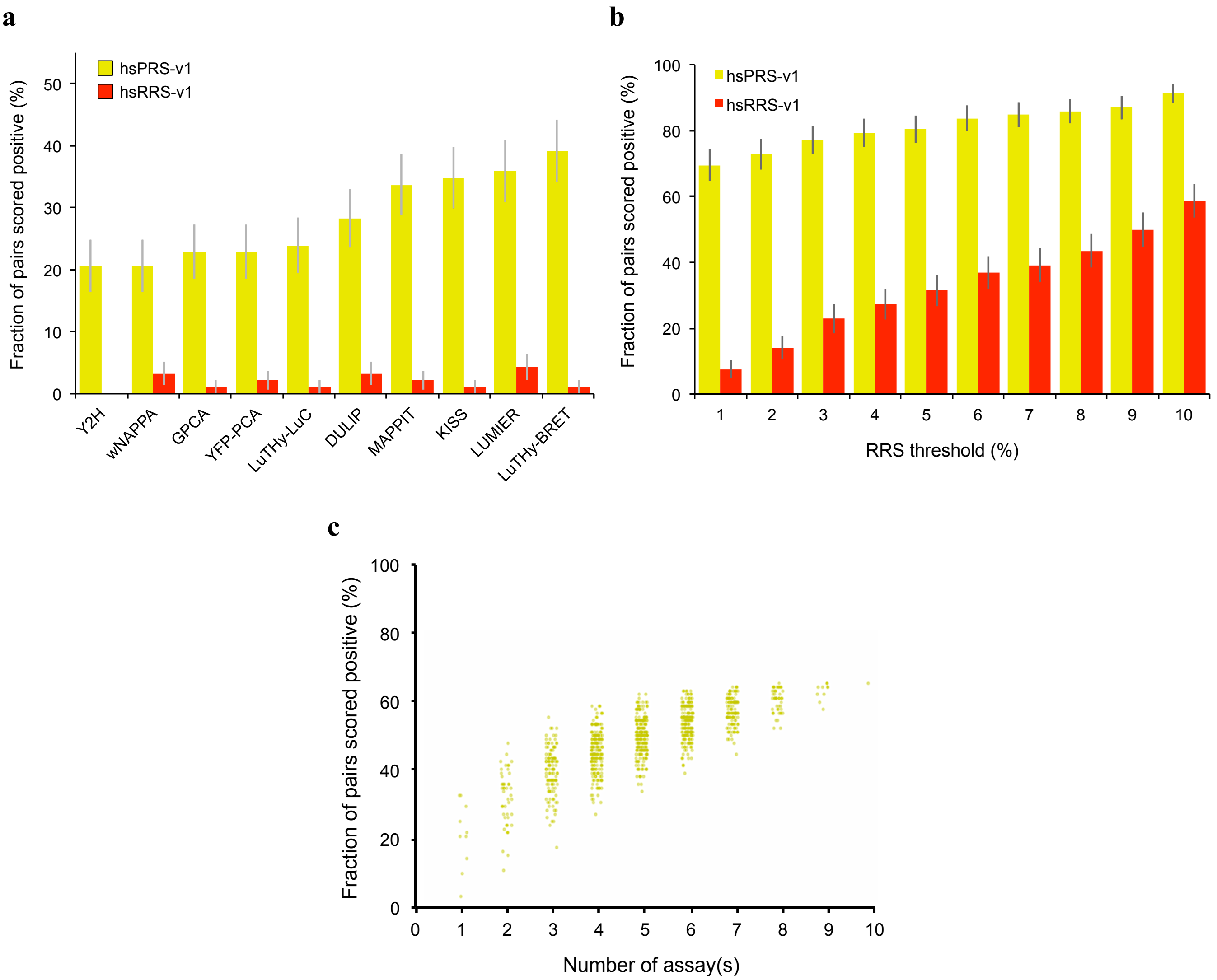


**Supplementary Figure 1 | Combining assays increases binary PPI detection.** (**a**) Recovery rates for ten binary PPI assays benchmarked against hsPRS-v1 and hsRRS-v1 as published by the authors in their original studies[1-6]. (**b**) Impact of scoring random pairs when combining assays: cumulative detection rates of hsPRS-v1 and hsRRS-v1 pairs when increasing, identical RRS thresholds are applied to YFP-PCA, LUMIER, MAPPIT, KISS, wNAPPA, LuTHy-BRET, LuTHy-LuC, GPCA and DULIP assays. (**c**) Cumulative hsPRS-v1 recovery rates when assays presented in **Fig. 2a** are combined: every possible combination of *n* assay(s) is displayed (yellow dots). See **Supplementary Table 1** for detailed description of assays. Error bars indicate standard errors of proportions (**a**) or standard deviations (**b**).

**a**

| Category | Protein (number of pairs involved) |
|---|---|
| PRS | TAB3, S100B(2), CCNA1(2), PTK2(2), PTPN11(2), RPA3, BNIP2, CD58, MAPK7, RAF1, CDK6, SMAD3 |
| RRS | FGF18, TRIM26, PSTPIP2, ATOH7, ENOX1, NUP37, TAB3, ZNF440, HNRNPM, JAK3, PML, PPP1R12B, HMGB1, PPP1R1A |

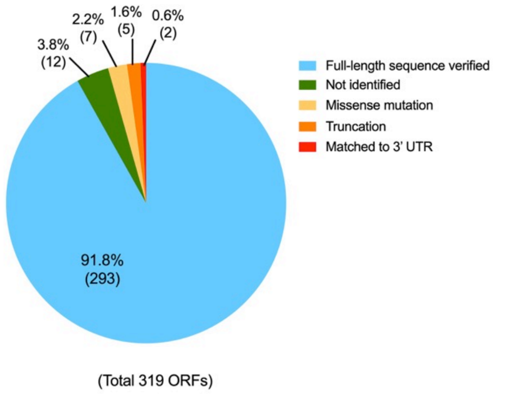


**b**

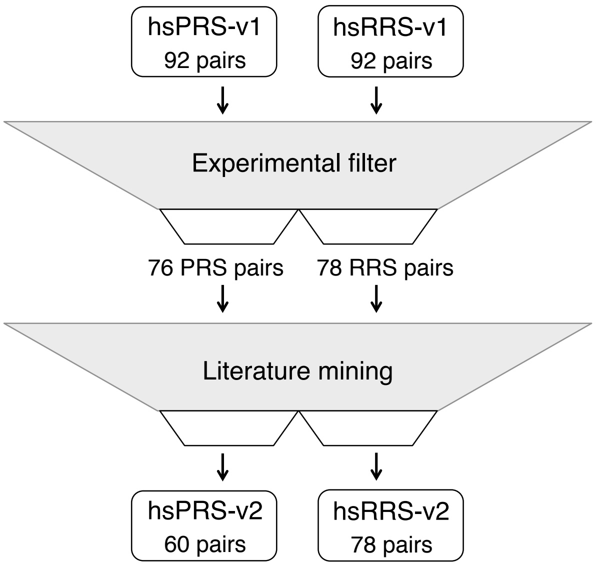


**c** **d**

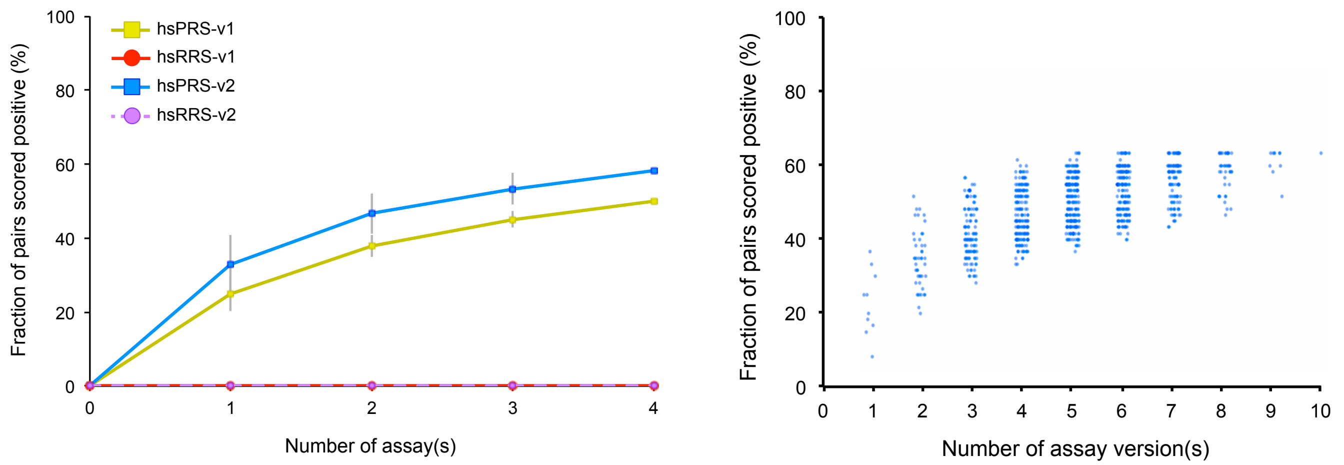


**Supplementary Figure 2 | Construction of second-generation positive and random reference sets.** (**a**) Classification and percentages of ORFs after experimental filtering of hsPRS-v1 and hsRRS-v1. Numbers in parentheses in the Table indicate the number of protein pairs in which the ORF is involved (if >1). (**b**) Summary of filtering steps applied to original hsPRS-v1 and hsRRS-v1 to construct second-generation hsPRS-v2 and hsRRS-v2. Experimental filter includes a series of ORF sequencing and clone purification, and literature mining includes filtering ORFs and protein pairs based on updated genome annotation and classification of binary PPI detection methods (see Methods for more details). Numbers of PRS and RRS pairs at each step are indicated. (**c**) Cumulative fractions of hsPRS-v1 and hsPRS-v2 recovered by Y2H (X-Y and Y-X versions), KISS (N1N2 and C1N2 versions), MAPPIT (X-Y and Y-X versions) and GPCA (N1N2 version) when none of the RRS pairs are scored positive. (**d**) Cumulative hsPRS-v2 recovery rates when assay versions presented in **Fig. 2c** are combined (excluding NanoBiT): every possible combination of *n* assay version(s) is displayed (blue dots). See **Supplementary Table 1** for detailed description of assays. Error bars in (**c**) indicate standard deviations.

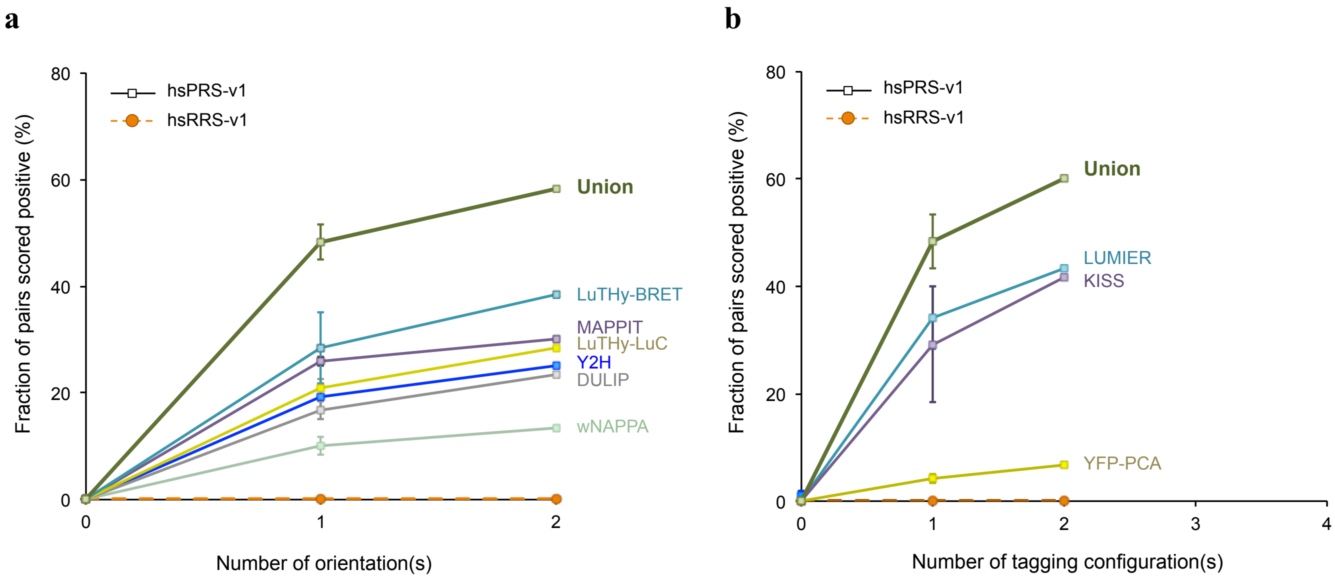


**Supplementary Figure 3 | Combination of published binary PPI assay versions.** (**a**), (**b**) PPIs detected by permuting (**a**) protein orientations in wNAPPA, DULIP, Y2H, LuTHy-BRET, LuTHy-LuC and MAPPIT, and (**b**) tagging configurations in LUMIER, YFP-PCA and KISS (only two versions tested for each of these assays; **Supplementary Table 1**). Assays were benchmarked with hsPRS-v1 and hsRRS-v1. Presented published results were re-analyzed and restricted to hsPRS-v2 and hsRRS-v2 spaces, at a threshold of no RRS pair scoring positive. Error bars indicate standard deviations.

a

b

c

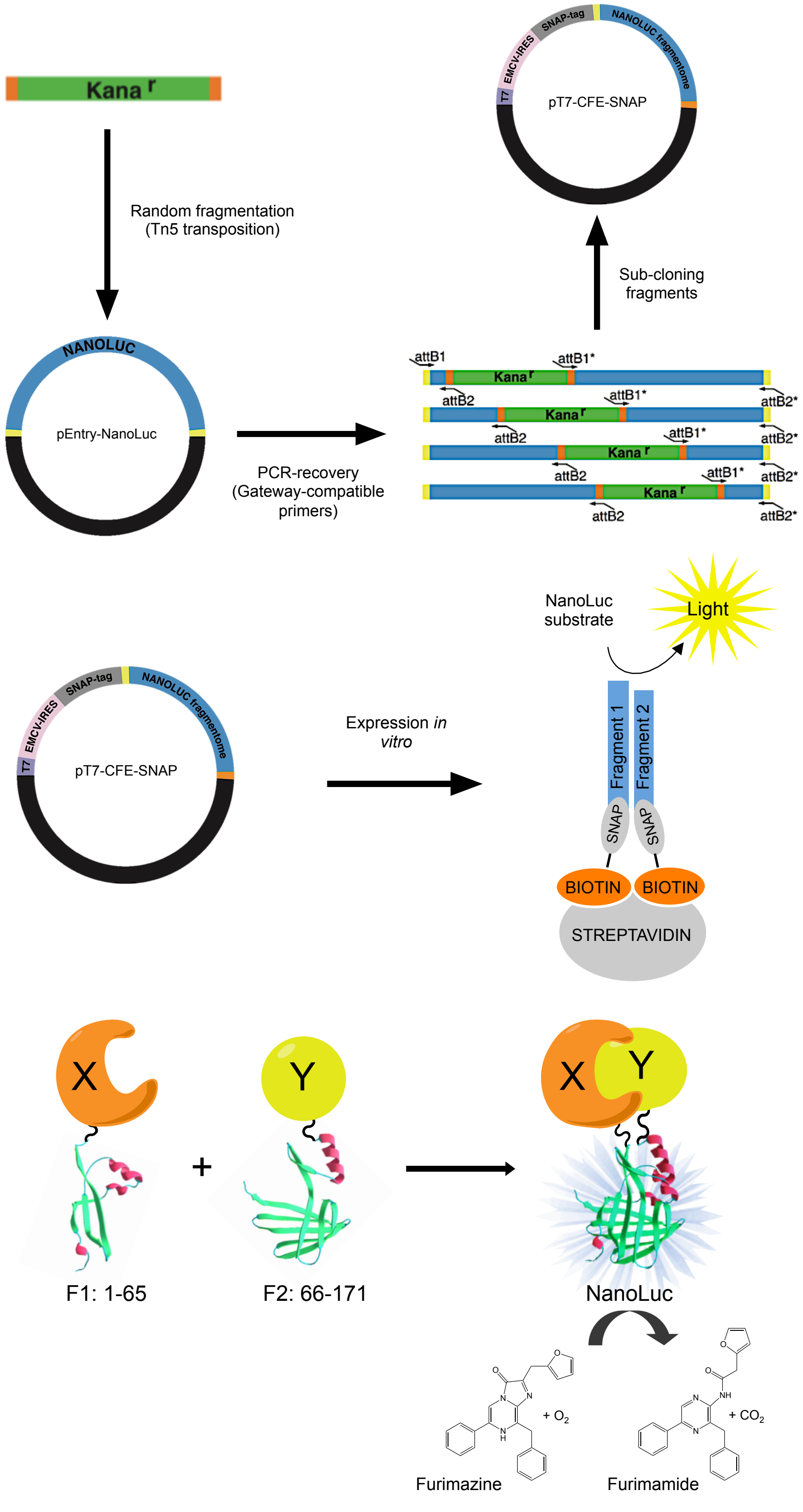


**Supplementary Figure 4 | Development of the N2H assay.** (**a**) NanoLuc fragmentation by transposon mediated random insertional mutagenesis. (**b**) *In vitro* proximity-based assay used to test complementation of fragments. (**c**) Principle of N2H assay when furimazine[7] is used as substrate.

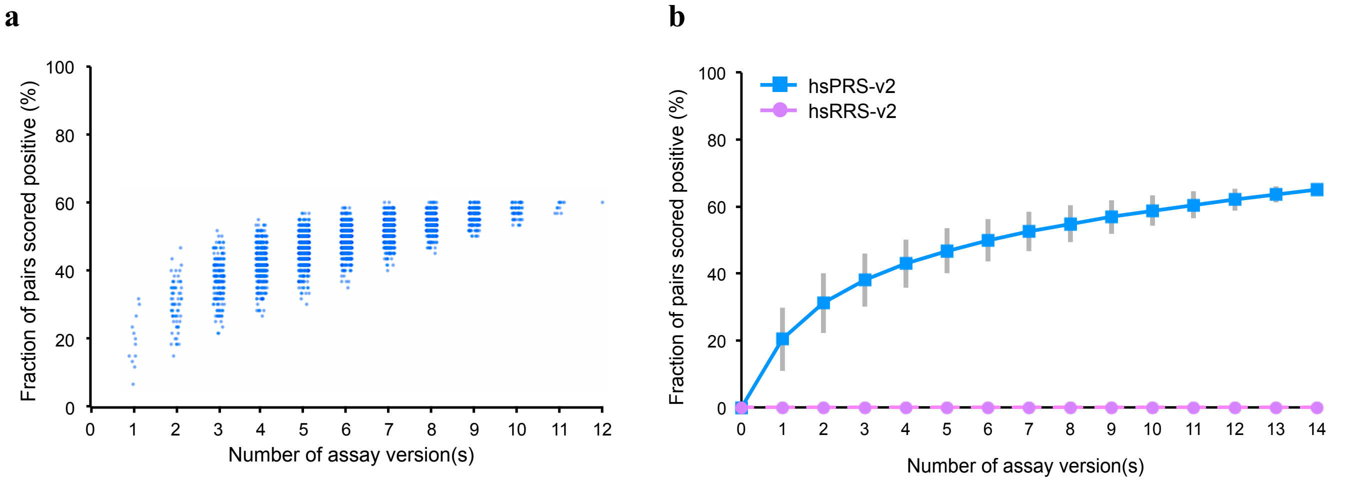


**Supplementary Figure 5 | Combination of binary PPI assay versions benchmarked against hsPRS-v2 and hsRRS-v2.** Cumulative hsPRS-v2 recovery rates when combining versions of (**a**) N2H presented in **Fig. 3e** (every possible combination of *n* assay version(s) is displayed (blue dots)), and (**b**) Y2H, MAPPIT, KISS, GPCA and NanoBiT presented in **Fig. 2c**. Error bars in (**b**) indicate standard deviations.

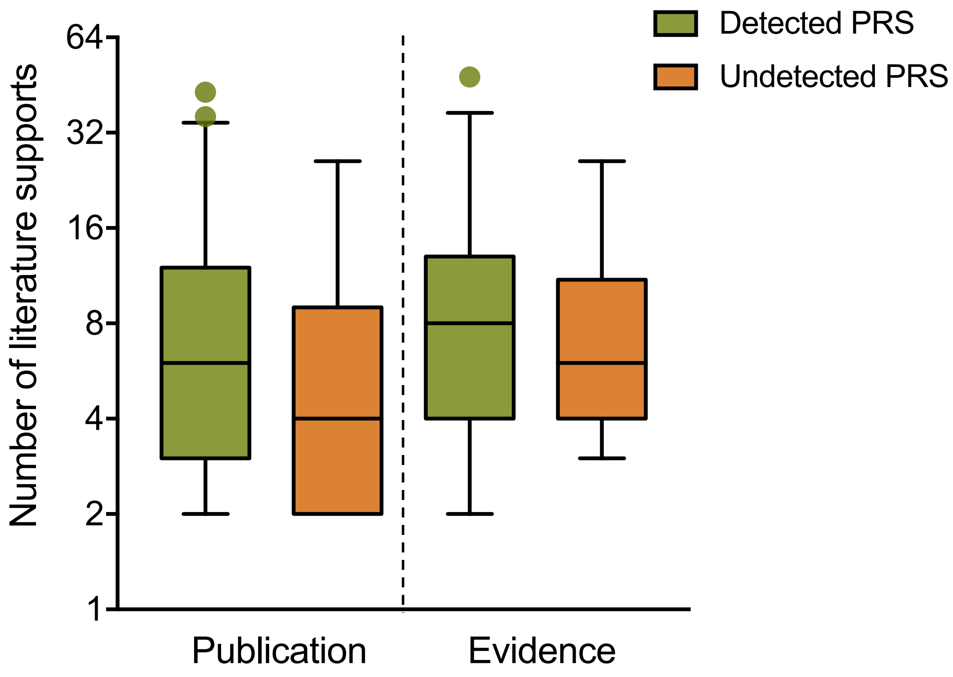


**Supplementary Figure 6 | Literature evidence for detected and undetected hsPRS-v2 pairs.** Box plots indicate the number of literature supports for detected (green) or undetected (orange) PRS pairs by all assays reported in **Fig. 6**. Left and right side graphs indicate the number of publications and the number of evidence, *i.e.* the combined number of publications and assays, for each PRS pair, respectively. Error bars show 5-95 percentile ranges.

### SUPPLEMENTARY TABLES

| Assay | Parameters tested with hsPRS- and hsRRS-v1 or v2 (this study) | | | | | Reference(s) |
|---|---|---|---|---|---|---|
| | Protein orientation | | Tagging configuration | | Expression environment | |
| | v1 | v2 | v1 | v2 | | |
| YFP-PCA | X-Y | n.t. | N1N2, N1C2 | n.t. | m | [1] |
| LUMIER | X-Y | n.t. | N1N2, N1C2 | n.t. | m | [1] |
| MAPPIT | X-Y/Y-X | X-Y/Y-X | N1N2 | N1N2 | m | [1] |
| KISS | X-Y | X-Y | N1N2, N1C2 | N1N2, N1C2 | m | [3] |
| GPCA | X-Y | X-Y | N1N2 | N1N2, N1C2, C1C2, C1N2 | m | [2], [8] |
| DULIP | X-Y/Y-X | n.t. | r.c. | n.t. | m | [4] |
| NanoBiT | n.t. | X-Y | n.t. | N1N2, N1C2, C1C2, C1N2 | m | [9] |
| LuTHy-BRET | X-Y/Y-X | n.t. | N1N2 | n.t. | m | [5] |
| LuTHy-LuC | X-Y/Y-X | n.t. | N1N2 | n.t. | m | [5] |
| Y2H | X-Y/Y-X | X-Y/Y-X | N1N2, N1C2, C1C2, C1N2 | N1N2 | y | [1], [6] |
| wNAPPA | X-Y/Y-X | n.t. | N1C2 | n.t. | v | [1] |
| N2H | | X-Y | | N1N2, N1C2, C1C2, C1N2 | m, y, v | This study |

n.t. = not tested; X, Y = tested proteins; r.c. = random configurations tested; y = yeast cells; m = mammalian cells; v = *in vitro* (cell-free)

**Supplementary Table 1 | Summary of assays and assay versions benchmarked against hsPRS-v1/hsRRS-v1 (v1; published data) or hsPRS-v2/hsRRS-v2 (v2; this study).** The Y2H versions highlighted in red were not used in this study, as non-ambiguous titrations of hsPRS-v1 pairs at different thresholds of hsRRS-v1 pairs scoring positive could not be obtained from the original study[6].

| Protein X | Protein Y | Reason | Details |
|---|---|---|---|
| GRAP2 | LAT | No evidence of interaction in binary PPI assay | Pull down, co-immunoprecipitation |
| NR3C1 | RELA | | Pull down, co-immunoprecipitation |
| PRKAR2A | EZR | | Co-immunoprecipitation, filter binding, affinity chromatography technology |
| LGALS3 | LGALS3BP | | Co-immunoprecipitation, fluorescence microscopy |
| AKT1 | TCL1A | | Pull down, co-immunoprecipitation |
| LCP2 | VAV1 | | Pull down, co-immunoprecipitation, affinity technology |
| HDAC1 | RB1 | | Pull down, co-immunoprecipitation, chromatography technology |
| RET | FRS2 | | Pull down, co-immunoprecipitation |
| FABP5 | S100A7 | | Co-immunoprecipitation |
| CRK | PDGFRB | | Pull down |
| B2M | HLA-A | No reference ORF | No consensus sequences for HLA proteins |
| B2M | HLA-C | | |
| B2M | HLA-B | | |
| FGF1 | FGFR1 | Proteolytic maturation of precursors | Precursor ligand/receptor protein |
| CXCL1 | CXCR2 | | |
| TNFSF10 | TNFRSF10B | | |

**Supplementary Table 2 | Bioinformatics and literature-based filters for discarded hsPRS-v1 pairs.**

| | Split-NanoLuc PPI detection technologies | | | |
|---|---|---|---|---|
| Assay name | NanoBiT | Unnamed | NanoPCA | N2H |
| Year, [reference] | 2015, [9] | 2015, [10] | 2017, [11] | This study |
| Number of fragment pairs tested | 90 | 9 | 7 | 58 |
| Modification of fragments? | Yes | No | No | No |
| F1 (N-terminal) | 11S (156 aa) | 1-52 | 1-67 | 1-65 |
| F2 (C-terminal) | Peptide 114 (11 aa) | 53-171 | 67-171 | 66-171 |
| Number of interaction(s) tested | 6 | 1 | 12 | 60 |
| Number of random protein pair(s) tested | 0 | 0 | 0 | 78 |
| Expression environment(s) | m | m | m | m, y, v |
| (F1:F2) / (full-length NLuc) Signal post-expression | n.r. | n.r. | n.r. | m: $1.3x10^{-6}$<br>y: $6.5x10^{-5}$<br>v: $3.1x10^{-6}$ |
| Comment | Steric constraints noticed when 11S on N-terminal | / | / | / |

aa = amino acid(s); m = mammalian cells; y = yeast cells; v = *in vitro* (cell-free); NLuc = NanoLuc; n.r. = not reported

**Supplementary Table 3 | Comparison of split-NanoLuc binary PPI detection technologies.** Different criteria are used for comparison. F1 and F2 represent the NanoLuc (NLuc) fragments used in the different split NanoLuc-based assays.

| | PROTEIN X | PROTEIN Y | PAIR # |
|---|---|---|---|
| hsPRS-v2 pairs | DR1 | DRAP1 | 1 |
| | LMNA | LMNB1 | 2 |
| | JUNB | BATF | 3 |
| | NCBP1 | NCBP2 | 4 |
| | LCP2 | GRAP2 | 5 |
| | BAK1 | BCL2L1 | 6 |
| | BAD | BCL2L1 | 7 |
| | SKP1 | BTRC | 8 |
| | SKP1 | SKP2 | 9 |
| | FANCA | FANCG | 10 |
| | PSMD4 | RAD23A | 11 |
| | LSM3 | LSM2 | 12 |
| | MAD2L1 | MAD1L1 | 13 |
| | PEX19 | PEX16 | 14 |
| | MAFG | NFE2L1 | 15 |
| | PEX14 | PEX19 | 16 |
| | PEX19 | PEX11B | 17 |
| | DDIT3 | FOS | 18 |
| | FEN1 | PCNA | 19 |
| | ATF3 | DDIT3 | 20 |
| | GTF2F1 | GTF2F2 | 21 |
| | IGF2 | IGFBP4 | 22 |
| | CDK2 | CKS1B | 23 |
| | PEX19 | PEX3 | 24 |
| | LCP2 | NCK1 | 25 |
| | CBLB | GRB2 | 26 |
| | MCM2 | MCM3 | 27 |
| | AKT1 | PDPK1 | 28 |
| | RCC1 | RAN | 29 |
| | NF2 | HGS | 30 |
| | TP53 | UBE2I | 31 |
| | HIF1A | TP53 | 32 |
| | GRB2 | VAV1 | 33 |
| | SMAD1 | SMAD4 | 34 |
| | CEBPG | FOS | 35 |
| | RHOA | ARHGAP1 | 36 |
| | SMAD4 | DCP1A | 37 |
| | CASP2 | CRADD | 38 |
| | XIAP | CASP9 | 39 |
| | NR3C1 | HSP90AA1 | 40 |
| | ORC2 | ORC4 | 41 |
| | RAC1 | ARFIP2 | 42 |
| | ERBB3 | NRG1 | 43 |
| | CGA | CGB5 | 44 |
| | ARF1 | ARFIP2 | 45 |
| | LMNA | RB1 | 46 |
| | XIAP | CASP7 | 47 |
| | IFIT1 | EIF3E | 48 |
| | ORC2 | MCM10 | 49 |
| | BDNF | NTF4 | 50 |
| | HDAC1 | ZBTB16 | 51 |
| | XIAP | CASP3 | 52 |
| | GADD45A | PCNA | 53 |
| | FANCA | FANCC | 54 |
| | RIPK2 | NOD1 | 55 |
| | GRB2 | LAT | 56 |
| | PDE4D | RACK1 | 57 |
| | PPP3CA | PPP3R1 | 58 |
| | MCM2 | MCM5 | 59 |
| | HBA2 | HBB | 60 |

| | PROTEIN X | PROTEIN Y | PAIR # |
|---|---|---|---|
| hsRRS-v2 pairs | ACVR1 | CWF19L1 | 61 |
| | APOD | MUC7 | 62 |
| | ARSA | DBN1 | 63 |
| | ASS1 | GINS3 | 64 |
| | ATP5O | CLEC2D | 65 |
| | BMP5 | C10orf119 | 66 |
| | BTC | BAT2L1 | 67 |
| | BYSL | KIAA0907 | 68 |
| | CA2 | PTPRS | 69 |
| | CANX | ARHGEF15 | 70 |
| | CANX | ATAD2 | 71 |
| | CD151 | WDR41 | 72 |
| | CD34 | SNX21 | 73 |
| | CD81 | NPC2 | 74 |
| | CENPA | PPIL3 | 75 |
| | CKB | HBZ | 76 |
| | CLPTM1 | L3MBTL2 | 77 |
| | CNN1 | PGAP2 | 78 |
| | COPB1 | HPCAL4 | 79 |
| | COPB1 | SLC39A14 | 80 |
| | DEFA3 | TSTD2 | 81 |
| | DLX4 | RAB3IP | 82 |
| | DUT | C19orf40 | 83 |
| | EMD | ARMC1 | 84 |
| | ERBB3 | C3orf38 | 85 |
| | ETF1 | LMBR1L | 86 |
| | FABP4 | GCG | 87 |
| | FABP7 | STX5 | 88 |
| | FAS | LSM3 | 89 |
| | FIGF | ZBTB25 | 90 |
| | FKBP3 | NQO2 | 91 |
| | GALK1 | MCCC1 | 92 |
| | GCDH | ZCCHC9 | 93 |
| | GP1BA | PVRL2 | 94 |
| | GPD2 | C22orf29 | 95 |
| | GPR18 | HNRPLL | 96 |
| | GRIK2 | ARL6IP6 | 97 |
| | HCLS1 | SALL2 | 98 |
| | HIST1H1C | NPDC1 | 99 |
| | HLA-DMB | PSEN2 | 100 |
| | INPP1 | UBLCP1 | 101 |
| | ITPA | WDR62 | 102 |
| | ITPK1 | TMEM22 | 103 |
| | LAMP2 | UBE2G2 | 104 |
| | LUM | UGGT2 | 105 |
| | MAOB | CTCF | 106 |
| | MCM2 | PLXNA4 | 107 |
| | MNAT1 | GMPPA | 108 |
| | MOBP | MRPS25 | 109 |
| | NAT2 | DNAJA1 | 110 |
| | NDP | NUDT4 | 111 |
| | NFIB | TIRAP | 112 |
| | NKX2-5 | CSGALNACT2 | 113 |
| | NONO | BEST1 | 114 |
| | NUDT2 | MIIP | 115 |
| | OSM | ZNF688 | 116 |
| | PBX2 | VILL | 117 |
| | PDE9A | REM2 | 118 |
| | PDGFRA | NDFIP1 | 119 |
| | PDHB | ZC3HC1 | 120 |
| | PMCH | RIC3 | 121 |
| | PPP6C | ZNF350 | 122 |
| | PROS1 | STK25 | 123 |
| | PSMD12 | CRIPT | 124 |
| | PSMD5 | SLC22A15 | 125 |
| | PSMD5 | SYCE1 | 126 |
| | RAB3B | BOC | 127 |
| | RBM3 | SF3A1 | 128 |
| | RCC1 | KLHL6 | 129 |
| | RFX3 | SEMA4G | 130 |
| | RGR | ABCF3 | 131 |
| | RHOC | NUP62CL | 132 |
| | RXRB | CXCL11 | 133 |
| | SCARB1 | PHF21B | 134 |
| | SERPINB3 | DCTN6 | 135 |
| | SHMT2 | STAC3 | 136 |
| | SLC25A6 | ZNF213 | 137 |
| | SLC6A1 | TM4SF4 | 138 |

**Supplementary Table 4 | hsPRS-v2 and hsRRS-v2 pairs.** Identity and given number for each pair of v2 sets (**Fig. 6**).

| Protein X | Protein Y | Pair number | Interaction domains present in tested ORFs? |
|---|---|---|---|
| IFIT1 | EIF3E | 48 | N.A. |
| ORC2 | MCM10 | 49 | N.A. |
| BDNF | NTF4 | 50 | N.A. |
| HDAC1 | ZBTB16 | 51 | N.A. |
| XIAP | CASP3 | 52 | Yes |
| GADD45A | PCNA | 53 | Yes |
| FANCA | FANCC | 54 | No, shorter isoform for FANCA |
| RIPK2 | NOD1 | 55 | Yes |
| GRB2 | LAT | 56 | Yes |
| PDE4D | RACK1 | 57 | No, shorter isoform for PDE4D |
| PPP3CA | PPP3R1 | 58 | Yes |
| MCM2 | MCM5 | 59 | Yes |
| HBA2 | HBB | 60 | Yes |

**Supplementary Table 5 | Undetected hsPRS-v2 pairs and analysis of published interaction domains.** Presence or absence of reported interaction domains in hsPRS-v2 ORF sequences. N.A.: not applicable (no domain reported); Yes: both ORFs contain reported domains. No: lack of reported interaction domain.

#### Supplementary references


1. Braun, P. et al. An experimentally derived confidence score for binary protein-protein interactions. *Nat. Methods* **6**, 91-97 (2009).
2. Cassonnet, P. et al. Benchmarking a luciferase complementation assay for detecting protein complexes. *Nat. Methods* **8**, 990-992 (2011).
3. Lievens, S. et al. Kinase substrate sensor (KISS), a mammalian in situ protein interaction sensor. *Mol. Cell. Proteomics* **13**, 3332-3342 (2014).
4. Trepte, P. et al. DULIP: a dual luminescence-based co-immunoprecipitation assay for interactome mapping in mammalian cells. *J. Mol. Biol.* **427**, 3375-3388 (2015).
5. Trepte, P. et al. LuTHy: a double-readout bioluminescence-based two-hybrid technology for quantitative mapping of protein-protein interactions in mammalian cells. *Mol. Syst. Biol.* **14**, e8071 (2018).
6. Chen, Y.C., Rajagopala, S.V., Stellberger, T. & Uetz, P. Exhaustive benchmarking of the yeast two-hybrid system. *Nat. Methods* **7**, 667-668 (2010).
7. Coutant, E.P. et al. Gram-scale synthesis of luciferins derived from coelenterazine and original insights into their bioluminescence properties. *Org. Biomol. Chem.* **10**, 3709-3713 (2019).
8. Sahni, N. et al. Widespread macromolecular interaction perturbations in human genetic disorders. *Cell* **161**, 647-660 (2015).
9. Dixon, A.S. et al. NanoLuc complementation reporter optimized for accurate measurement of protein interactions in cells. *ACS Chem. Biol.* **11**, 400-408 (2016).
10. Verhoef, L.G., Mattioli, M., Ricci, F., Li, Y.C. & Wade, M. Multiplex detection of protein-protein interactions using a next generation luciferase reporter. *Biochim. Biophys. Acta* **1863**, 284-292 (2016).
11. Mo, X. et al. AKT1, LKB1, and YAP1 revealed as MYC interactors with NanoLuc-based protein-fragment complementation assay. *Mol. Pharmacol.* **91**, 339-347 (2017).
